# Direct RNA Sequencing Reveals Stress-Dependent and Pathway-Specific rRNA Modification Reprogramming During 50S Biogenesis

**DOI:** 10.64898/2026.01.02.697427

**Authors:** Luis A. Gracia Mazuca, Salvador A. Rodarte, Isaac S. Weislow, Jonathon E. Mohl, Eda Koculi

## Abstract

Ribosomal RNA (rRNA) modification and processing are essential steps in ribosome assembly. Using Oxford Nanopore direct RNA sequencing, we simultaneously detect and quantify eight classes of 23S rRNA modifications in the mature 50S large subunit (LSU) from *Escherichia coli* cells expressing either wild-type DbpA or the helicase-inactive R331A DbpA variant, as well as in two LSU assembly intermediates, 35S and 45S, which accumulate along distinct maturation pathways in R331A DbpA expressing cells. In addition, we analyze 3′-end processing of 23S and 5S rRNAs across these particles. Many 23S rRNA modifications are incorporated at similar levels in LSU assembly intermediates and mature 50S subunits from both wild-type and R331A DbpA expressing cells, indicating that these modifications are incorporated prior to intermediate accumulation and are not preferentially reprogrammed under R331A DbpA induced assembly stress. In contrast, a subset of three modifications exhibits altered incorporation patterns. N^2^-methyladenosine 2507 incorporation is reduced in the 50S LSU from R331A DbpA expressing cells compared with the cells expressing wild-type DbpA, whereas pseudouridine (Ψ) 2508 is increased. In addition, Ψ 2608 is reduced in the 50S subunit from R331A DbpA expressing cells compared with the 35S and 45S intermediates from the same cells and the 50S subunit from wild-type cells. Because the 35S and 45S pathways account for only ∼40% of ribosome assembly in R331A DbpA expressing cells, these findings demonstrate that Ψ2608 incorporation is selectively reprogrammed across alternative *in vivo* assembly routes, revealing an additional regulatory layer in ribosome biogenesis.

## INTRODUCTION

RNA post-transcriptional modifications are ubiquitous in all organisms, regulating RNA structure, RNA-protein interactions, RNA transport, and RNA fate ^1–7^. Furthermore, several enzymes responsible for installing these modifications function as RNA chaperones ^8–11^. After transfer RNA (tRNA), the ribosomal RNA (rRNA) molecule is the most heavily modified RNA molecule in the cell ^6^. In bacteria, the extent of rRNA modifications and when they are incorporated during ribosome assembly have been shown to shift in response to metabolic and oxidative stresses, cold shock, physiological changes, and antibiotic exposure ^12–17^. Here, we investigate how eight distinct classes of rRNA modifications are reprogrammed during impaired ribosome assembly caused by the expression in the *Escherichia coli* (*E. coli*) cells of helicase inactive R331A DbpA protein ^18^. Understanding this reprogramming provides novel biochemical insight into the dynamic regulation of ribosome biogenesis and underscores the role of RNA modifications in preserving ribosome functionality during cellular adaptation to physiological stress.

The DbpA protein is a DEAD-box RNA helicase that performs structural rearrangements in the peptidyl transferase center (PTC) of the large ribosomal subunit (LSU) (50S) ^19–26^. The DbpA protein does not interact with the small subunit (SSU) (30S) ^19, 27^. The expression of R331A DbpA produces slow growth and a cold-sensitive phenotype and leads to the accumulation of three large subunit intermediates with sedimentation coefficients of 27S, 35S, and 45S ^18, 28, 29^. The three intermediates rearrange to form 50S LSU via independent pathways^28^.

LSU assembly involves extensive rRNA folding and rearrangement, and chemical modifications have been shown to facilitate maturation of the 50S subunit ^7, 10, 11, 30^. The 50S LSU contains two RNA molecules, 5S rRNA and 23S rRNA ^31^. While 5S rRNA does not harbor known RNA modifications, 23S rRNA contains 25 chemically modified nucleotides (Table S1)^16, 27, 30^. Our previous studies using Illumina next-generation sequencing (NGS) characterized the incorporation of 17 modifications (1-methylguanosine 747 (m^1^G 747), pseudouridine 748 (Ψ 748), Ψ 957, N^2^-methylguanosine 1837 (m^2^G 1837), Ψ 1915, 3-methylpseudouridine 1919 (m^3^Ψ 1919), Ψ 1921, 7-methylguanosine 2073 (m^7^G 2073), m^2^G 2449, dihydrouridine 2453 (D 2453), Ψ 2461, 5-hydroxycytidine 2505 (OH^5^C 2505), N^2^-methyladenosine 2507 (m^2^A 2507), Ψ 2508, Ψ 2584, Ψ 2608, and Ψ 2609) in 23S rRNA from the 27S, 35S, and 45S LSU assembly intermediates accumulating in the cells expressing R331A DbpA protein and compared their levels with those in the mature 50S subunit from cells expressing wild-type DbpA^32–34^. These analyses established whether the modifications were incorporated into assembly intermediates to the same extent as in the mature LSU from the cells expressing wild-type DbpA ^32–34^.

However, these studies did not address whether modification levels differ between mature 50S subunits isolated from cells expressing wild-type DbpA and those expressing the helicase-inactive R331A DbpA construct ^32–34^. Addressing this question is important because comparing modification levels in mature 50S subunits from wild-type and R331A DbpA expressing cells directly tests whether R331A DbpA induced ribosome assembly stress reprograms the final rRNA modification landscape.

In addition, eight 23S rRNA modifications (N^6^-methyladenosine 1620 (m^6^A 1620), 5-methyluridine 1943 (m^5^U 1943), 5-methylcytidine 1966 (m^5^C 1966), m^6^A 2034, 2′-O- methylguanosine 2255 (G_m_ 2255), 2′-O-methylcytidine 2502 (C_m_ 2502), and 2′-O-methyluridine 2556 (U_m_ 2556)) were not accessible using Illumina NGS in either LSU assembly intermediates or the mature 50S subunit, excluding assessment of their incorporation timing and regulation during assembly in the R331A DbpA expressing cells ^32–35^.

To overcome these limitations, we employed Oxford Nanopore direct RNA sequencing (ONT DRS) to analyze modification incorporation in the 35S and 45S LSU assembly intermediates that accumulate in cells expressing R331A DbpA, and to directly compare modification extents in mature 50S subunits isolated from cells expressing either wild-type DbpA or the R331A DbpA variant. This approach enabled analysis of several modifications previously characterized by Illumina NGS and additionally allowed detection of m^5^ U749 and U_m_ 2556, which were not detectable using Illumina NGS ^32–34^. Collectively, the ONT DRS data presented here allow us to determine whether selective rRNA modifications are reprogrammed under R331A DbpA induced LSU assembly stress, both in the mature 50S subunit and across distinct *in vivo* assembly pathways populated in R331A DbpA expressing cells.

## MATERIALS AND METHODS

### Materials

ONT DRS kits (SQK-RNA002), Flow Cell Priming Kits (EXP-FLP002), Flow Cell Wash Kits (EXP-WSH004), and Flow cells were obtained from Oxford Nanopore Technologies. *E. Coli* RNA Poly(A) Polymerase, 10X *E. Coli* Poly(A) Polymerase Reaction Buffer, 10 mM ATP, T4 DNA Ligase, 10 mM dNTP Solution, and Quick Ligation Reaction Buffer, DNase I, RNase-free, were obtained from New England Biolabs (NEB). Dichlorodiphenyltrichloroethane (DDT), SuperScript III Reverse Transcriptase, Qubit RNA HS Assay Kits, and Qubit 1x dsDNA HS Assay Kits, MEGAscript^TM^ T7 Transcript Kit were obtained from ThermoFisher. Agencourt Ampure XP Reagent was obtained from Beckman Coulter, RNasin was purchased from Promega. The RNeasy kit was purchased from Qiagen. NaCl, Tryptone, Yeast extract, HEPES, NH_4_Cl, MgCl_2_, sucrose and KOH were purchased from Fisher Scientific. 2-mercaptoethanol (BME) and Lysozyme were purchased from Sigma-Aldrich.

### Isolations of ribosomal particles and rRNA molecules

The ribosomal particles were isolated from the same cells and in a similar manner as in our previous studies, with the exception that BioComp system combined with the Gilson fraction collector were used to fractionate the sucrose gradient and collect the fractions instead of the Teledyne R1 Fraction Collector and Brandel SYR-101 Syringe Pump system ^32, 33^. In brief, ribosomal particles were isolated from *E. coli* BLR (DE3) *plys*S, Δ*dbpA*/*kanR* cells expressing the wild-type DbpA or R331A DbpA ^32, 33^. The DbpA constructs were expressed from pET-3a vector in absence of IPTG. Thus, these constructs were produced from the leaky expression of the pET-3a vector ^32, 33^. The cell pellets were resuspended in a buffer consisting of 4mM BME, 200 mM HEPES, 30 mM NH_4_Cl, 1 mM MgCl_2_ and lysozyme 1mg/ml. The cells were broken by using multiple cycles of freeze-thaw. Next the DNA was digested by incubating the broken cells in ice with DNase I, RNase-free. After cell breakage and DNA digest, the cellular debris were separated from the lysate by centrifugation. The lysate was loaded on a linear 20 to 40% (weigh by volume) linear sucrose gradient made in buffer consisting of 4 mM BME, 50 mM HEPES pH 7.5, 150 mM NH_4_Cl, 1 mM MgCl_2_. To separate the ribosomal particles, the gradients were centrifuged at 126522 xg for 16.5 hours and subsequently fractionated using the BioComp system combined with the Gilson fraction collector.

The rRNA was isolated from the ribosomal particles by employing the RNeasy kit and followed the manufacturer’s directions.

### rRNA *in-vitro* transcription

The 23S rRNA was expressed from pCW1 plasmid. This plasmid was a gift from Olke Uhlenbeck’s laboratory at Northwestern University ^19^ . The DNA coding sequence of 16S rRNA was inserted into the KpnI and XbaI restriction enzyme sites of pUC19. The pUC19 with the 16S rRNA sequence was synthesized by GenScript. The pCW1 plasmid was linearized using AFLII restriction enzyme, while the plasmid carrying the 16S rRNA sequence was linearized using XbaI restriction enzyme. The complete linearization of the plasmid was confirmed using 1% agarose gel electrophoresis.

The 16S and 23S rRNA molecules were transcribed and isolated using the MEGAscript^TM^ T7 Transcript Kit following the manufacturer’s directions with the sole exception that 1 μl of RNase Inhibitor (40 u/mL) was added on the initial reaction mixture.

### rRNA and Nanopore Library Preparation

Prior to polyadenylation reaction, the RNA was denatured at 90°C in water for two minutes and immediately placed in ice to prevent structural formation. Polyadenylation of RNA was performed using the NEB Poly(A) polymerase protocol with minor modifications listed here. First, 0.5 mL RNase inhibitor (40 u/mL) was added to the polyadenylation reaction. Second, the reaction was allowed to proceed at 37°C for 30 minutes instead of the manufacturer’s suggested 15 minutes. Lastly, the polyadenylation reaction was not quenched by Ethylenediaminetetraacetic Acid (EDTA), instead this reaction was terminated by removing the Poly(A) polymerase and ATP with Agencourt AMPure XP Beads.

The determination of the rRNA concentrations used for the library preparation was obtained using Qubit Fluorometric Quantification. The library preparations were performed following the directions of ONT DRS Kit, SQK-RNA002, with three minor modifications listed here. First, before the reverse transcriptase adaptor ligation step, the rRNA was incubated at 90°C in water for 2 minutes and then immediately placed in ice. This step was not in the ONT protocol. The buffer, the adaptors and the T4 DNA ligase were added while the RNA was still in ice. Second, the above adaptor ligation mix was incubated at 22°C for 15 minutes, which was 5 minutes longer than the incubation time suggested by the ONT. Lastly, the mRNA provided by the ONT as a positive control for the nanopore library preparations were not employed in our study.

### Nanopore data analysis workflow

Basecalling was done using Guppy, a software provided by ONT, version 6.4.2+97a7f06, from ONT Community (Figure S1). This was the GPU version set up with the high-accuracy option for the RNA sequencing kit 9.4.1, as specified in the configuration file rna_r9.4.1_70bps_hac.cfg. The Guppy default settings were used, with a few minor modifications listed below. To put all basecalled reads into one fastq file, we set the parameter “ --records_per_fastq” to 0. We used the GPU to accelerate base calling in Guppy. Thus, we set the “--device” option in Guppy to cuda: all:100%.

For aligning sequences, we used the “--align_ref” option. We provided the reference RNA sequence to Guppy. Because we were interested in investigating the 3’ end processing of 5S and 23S rRNA molecules, we aligned the 5S and 23S rRNA to sequences that contained 15, and 20 extra nucleotides, respectively, in their 3’ ends. The sequence was part of the non-processed rRNA sequences of the *E. coli* genome.

Post-basecalling, FASTQ files, were merged and aligned using Minimap2 to *rrlB* gene or *rrsB* gene ^36^. The *rrlB* gene is one of the seven *E. coli* genes that encode 23S rRNA, and the *rrsB* gene is one of the seven genes that encode 16S rRNA. Next, low quality score reads were filtered out using SAMtools ^37^. As suggested by ONT, the reads with a quality score lower or equal than 10 were disregarded. A dictionary for reference genes was built using the Picard tool ^38^. This reference dictionary was required for alignment optimization by EpiNano, the program we employed to determine the RNA modifications ^39^. Subsequently, the BAM file was indexed using SAMtools ^37^.

The indexed BAM files were used by EpiNano (version 1.2.2) to obtain Z scores and the sum of errors (SE) ^39^. We combined the Z scores and the sum of errors to detect modifications and determine their extent in various ribosomal particles.

To determine the extents of correctly processed and unprocessed 5S and 23S rRNA molecules at their 3’ end, first we needed to extract the read depths for 3’end positions of 5S and 23S rRNA. The read depths were extracted by running SAMtools mpileup function ^37^. SAMtools mpileup function requires the BAM file obtained from Guppy’s built-in Minimap2 and the reference FASTA file ^36, 37^. The SAMTool mpileup function generated a table in CSV format which contained the read depths for each position in the sequence ^37^.

The number of reads for 5S rRNA was significantly smaller than the number of reads for the 16S and 23S rRNA. The above is consequence of two factors. First, the 5S rRNA is only 120 nucleotide long, during the library preparation steps, an extensive amount of 5S rRNA was lost^40^. Second, because of its small size, a number of 5S rRNA reads were counted as adaptors during the Guppy base calling process. To increase the number of 5S reads, we didn’t filter our FASTQ reads based on quality scores during Guppy base calling, as we did for the 16S and 23S rRNA. Thus, for the 5S rRNA base calling with Guppy we used the “--disable_qscore_filtering” command. Filtering out low quality data is important for the correct determination of errors, insertions and deletions required for detection of 16S and 23S rRNA modifications, as explained below. In the case of 5S rRNA, we were not concerned with RNA modifications, but only with the 3’ end processing.

### 16S rRNA and 23S rRNA modification detections

We adapted two outputs of EpiNano, the Z score and SE to quantify RNA modifications in ribosomal particles ^39^. The SE is computed by EpiNano as the addition of the deletions, insertions and mismatches at a position. EpiNano computed the sum of errors for our biological samples (SEB) and *in vitro* transcribed rRNA molecules (SEI).

The difference of the SEB between the biological sample and SEI was calculated by EpiNano using Equation 1^39^. In this equation, *i* is the nucleotide number position in 16S or 23S rRNA.

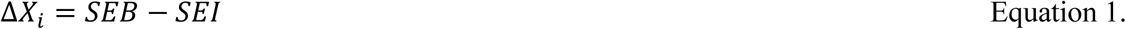

The mathematical mean, μ, was calculated using Equation 2 ^39^. In this equation, *n* is the last position in the sequences of the 16S or 23S rRNA molecules, and *i* and Δ*X_i_* are defined in Equation 1.

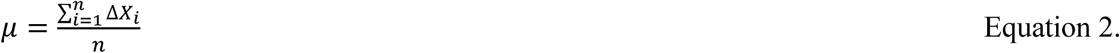

The standard deviation for the complete 16S or 23S rRNA sequencing of a biological replicate was calculated using Equation 3. The *i*, *n*, Δ*X_i_* and μ are defined in Equations 1 and 2.

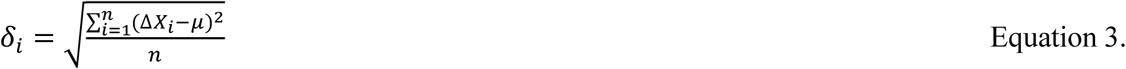

Z score for every base in the sequence was computed using the equation below, where Δ*X_i_* , μ and *δ _i_* are described in Equations 1-3 ^39^.

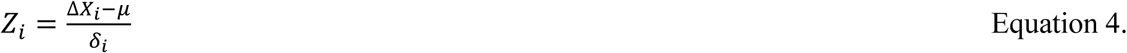

Lastly, we computed the ratios of SEB and SEI at respective nucleotides positions. A nucleotide is deemed modified if and only if it satisfies two thresholds: (I) the Z*_i_* value is ≥ 3, and ratio of SEB over SEI is ≥ 2.

### Determination of 3’ end 23S rRNA and 5S rRNA processing

The last steps of 3’ end 23S rRNA processing consists of constructs containing extra five and eight nucleotides. The correctly processed rRNA of *rrlB* gene is 2908 nucleotide long. The rRNA molecules with 5 extra nucleotides are 2913 nucleotides long, and the rRNA molecules with eight extra nucleotides are 2916 ^41^. As described in more details below, the fractions of different extents of processing were calculated by first determining the read depths of each 23S rRNA processed class and then dividing it by the total read depth for the 3’ end region.

To calculate the average number of mature 23S rRNA in different particles, we subtracted the read depth at position 2909 of 23S rRNA from the read depth of positions 2901 to 2908 of 23S rRNA. The position 2909 is the first unprocessed nucleotides. Next, we averaged the number of correctly processed reads for the residues 2901 to 2908. The same process was repeated for the 23S rRNA molecules with five extra bases. To determine the average number of 23S rRNA with five extra bases, the read depth at position 2914 was subtracted from the read depths of positions 2909 to 2913, and then these adjusted read depths were averaged. Lastly, the average read depth for the 23S rRNA molecules with eight extra nucleotide was calculated from the average of read depths of residues 2914 to 2916. Once the average read depths were calculated for the 23S rRNA molecules correctly processed, containing extra five or eight nucleotides, these averages were added to produce the total number of reads. The fractions of correctly processed 23S rRNA, the unprocessed rRNA containing five or eight extra bases were calculated by dividing the average read depth of each 23S rRNA processing class by the total number of reads.

The last steps of 3’ end 5S rRNA processing consists of molecules with three or more extra bases ^42^. The mature 5S rRNA is 120 nucleotides long. Our nanopore data show that for large subunit ribosomal intermediates isolated under our cellular conditions, the unprocessed 5S rRNA at 3’ end contained five extra bases. The fractions of 5S rRNA molecules, which were correctly processed or contained 5 extra bases at their 3’ end, were calculated as explained above for the 23S rRNA molecules. First, the read depth at position 121 was subtracted from the read depth in position 116 to 120. Then, the average of the adjusted read depths of positions 116 to 120 was calculated. Next, we calculated the average read depths of unprocessed 3’ end 5S rRNA, this consisted of positions 121 to 125. Lastly, the fractions of correctly processed and unprocessed 5S rRNA were calculated by dividing the adjusted average read depth of positions 116 to 120 and the adjusted average read depth of positions 121 to 125, respectively, by the sum of both averages.

## RESULTS AND DISCUSSION

### EpiNano can detect 16 out of 25 LSU rRNA modifications and 9 out of 11 SSU rRNA modifications

The post-transcriptionally modified 16S rRNA was isolated from the 30S subunit accumulating in cells expressing wild-type DbpA. The post-transcriptionally modified 23S rRNA was isolated from the 35S and 45S intermediates accumulating in cells expressing R331A DbpA, and from the 50S subunit in cells expressing either R331A or wild-type DbpA ^28, 32–34^. *In vitro* transcribed 16S and 23S rRNA, which contained no post-transcriptional modifications, were used to distinguish between background errors and those arising from modifications during ONT DRS sequencing, as explained in the Materials and Methods section of the manuscript.

Taking advantage of ONT DRS combined with EpiNano, we were able to detect 12 out of 17 classes of RNA modifications present in the 30S and 50S ribosomal subunits (Figure 1, Table S1, Table S2-S4) ^39^. A residue was deemed modified if and only if its Z-score, as calculated by EpiNano, was greater than or equal to 3, and the ratio of the sum of errors in RNA isolated from cells versus in *vitro*-transcribed control RNA was greater than or equal to 2 (Equation 4)^39^. Employing these thresholds, we were able to detect the following modifications in 23S rRNA from the 50S LSU isolated from cells expressing wild-type DbpA: m^1^G 747, Ψ 748, m^5^U 749, Ψ 957, Ψ 1915, m^3^Ψ 1919, Ψ 1921, m^7^G 2073, D 2453, Ψ 2461, m^2^A 2507, Ψ 2508, U_m_ 2556, Ψ 2584, Ψ 2608, and Ψ 2609 (Figure 1A). However, we were unable to detect in 23S rRNA isolated from 50S LSU accumulated in cells expressing wild-type DbpA the following RNA modifications: m^6^A 1620, m^2^G 1837, m^5^U 1943, m^5^C 1966, m^2^G 2449, G_m_ 2255, C_m_ 2502, OH^5^C 2505 (Figure 1A). A large number of modifications (m^1^G 747, Ψ 748, Ψ 957, Ψ 1915, m^3^Ψ 1919, Ψ 1921, m^7^G 2073, D 2453, Ψ 2461, m^2^A 2507, Ψ 2508, Ψ 2584, Ψ 2608) can be detected by both Illumina NGS and ONT DRS ^32–35^. On the other hand, m^5^U 749 and U_m_ 2556 are uniquely detected by ONT DRS (Figure 1A, Table S2, Table S4).

**Figure 1.**
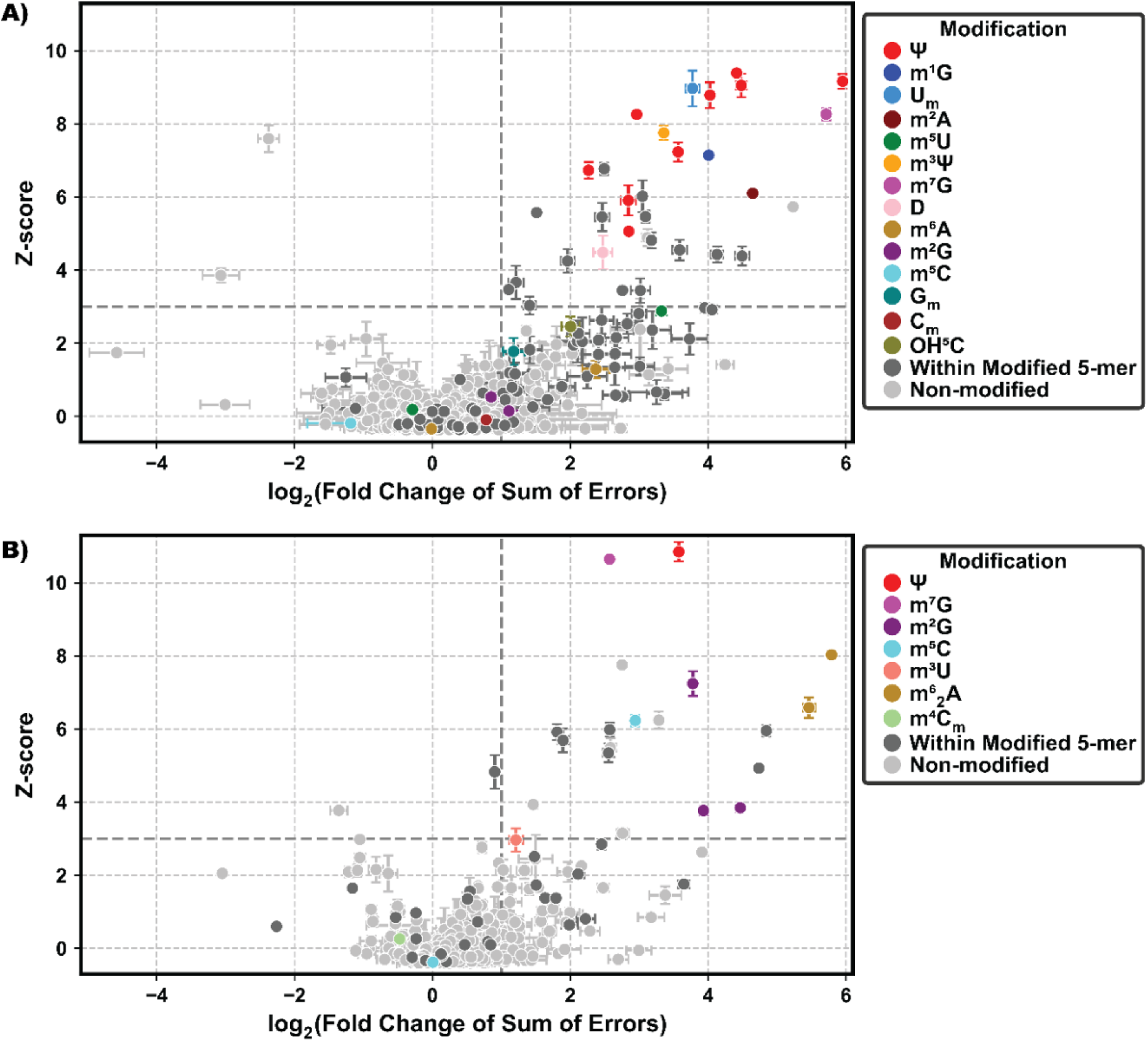
Oxford Nanopore sequencing combined with EpiNano detects 16 out of 25 rRNA modifications present in the 23S rRNA of the large ribosomal subunit and 9 out of 11 modifications present in the 16S rRNA of the small subunit ^39^. A) Modifications detected in 23S rRNA from the 50S large subunit isolated from wild-type cells. B) Modifications detected in 16S rRNA from the 30S small subunit isolated from wild-type cells. For both panels (A) and (B), the values shown represent averages obtained from two biological replicates. Error bars indicate the standard deviation of both the Z-score and fold-change values. As explained in the Materials and Methods section, only modifications with a Z-score ≥ 3 and a fold change ≥ 2 are considered detectable in this study. Z-scores and fold-change values for the large and small ribosomal subunit particles analyzed are available in Table S2, Table S3, Table S4.

In the SSU 16S rRNA we detected the following modifications: Ψ 516, m^7^G 527, m^2^G 966, m^5^C 967, m^2^G 1207, m^3^U 1498, m^2^G 1516, m^6^ A 1518, and m^6^ A 1519 (Figure 1B, Table S3, Table S4). However, in this RNA molecule we were unable to detect m^4^C_m_ 1402 and m^5^C 1407 modification (Figure 1B, Table S3, Table S4). Comparison of the modifications we detected in 23S rRNA and 16S rRNA reveals three classes of modification, m^5^U, m^5^C and m^2^G, are above our detection threshold in certain rRNA sequence but below it in others (Figure 1, Table S2, Table S3, Table S4). Thus, our detection of RNA modification using ONT DRS combined with EpiNano is sequence context dependent. This was also observed in previous ONT DRS studies^43^.

The data in Figure 1 show several nucleotides known to be unmodified yet displaying a Z-score greater than 3 and a fold change in the SEB-to-SEI ratio greater than 2. The vast majority of these nucleotides are located within three nucleotides of a modified site. EpiNano detects ONT DRS associated errors, insertions, deletions, and mismatches, by analyzing changes in ionic current ^39^. Thus, the data in Figure 1 reveal that current changes caused by RNA modifications at a given nucleotide can influence the ionic signal of adjacent nucleotides. This phenomenon has been previously observed in ONT DRS experiments ^12, 44^. Because several unmodified nucleotides exceed our detection thresholds, the ONT DRS combined with the EpiNano approach described here is not suitable for the discovery of novel RNA modifications; accordingly, identification of novel modifications was not a goal of this study (Figure A) ^39^. However, this approach enables robust quantification of known RNA modifications and their relative changes across samples, which is demonstrated by the strong agreement between two biological replicates in both the SE and ΔXᵢ values across all investigated samples (Figure S2-S7).

### m^1^G 747 and m^7^G 2073 are incorporated prior to 35S and 45S accumulation and remain unaffected in the 50S LUS under R331A DbpA induced assembly stress

The RlmA enzyme, which incorporates m^1^G 747, is located in the loop of helix 35 of 23S rRNA, near the entrance of the nascent peptide exit tunnel ^31, 45, 46^. Deletion of the *rlmA* gene produces a slow growth phenotype, decreased translation rate, and increase resistance to viomycin ^45^. Thus, investigating m^1^G 747 incorporation examines whether a modification implicated in translation efficiency, antibiotic resistance, and cell growth is reprogrammed during perturbed LSU assembly induced by R331A DbpA expression ^45^.

The RlmKL enzyme incorporates the m^7^G 2073 modification ^10^. This enzyme consists of two domains with distinct and independent functions ^10^. The K domain catalyzes methylation at the N7 position of G 2073, whereas the L domain catalyzes methylation at the C2 position of G 2449 ^10^. Both nucleotides, G 2073 and G 2449, are located within helix 74 of the PTC. *In vitro* studies using a fragment of 23S rRNA reveal that unwinding of helix 74 is required for the methyltransferase activity of both the K and L domains and that RlmKL facilitates this unwinding ^10^. Moreover, in cells lacking the DEAD-box RNA helicase DeaD, which is involved in LSU assembly, RlmKL has been shown to compensate for the loss of DeaD function ^10^. Recently, the RlmL domain was found to act as an RNA chaperone, promoting large subunit ribosome assembly independently of its methyltransferase activity ^11^. Given the role of RlmKL as an RNA chaperone that facilitates helix 74 remodeling during LSU maturation and installs the m^7^G 2073 modification within this helix, examining whether m^7^G 2073 incorporation is altered under R331A DbpA induced ribosome assembly stress tests whether this modification is sensitive to impaired PTC associated rRNA remodeling.

m^1^G 747 and m^7^G 2073 are incorporated at similar levels in the 35S, 45S, and 50S particles accumulating in cells expressing R331A DbpA, as well as in the 50S subunit from cells expressing wild-type DbpA (Figure 2, Table S2, Table S4). This indicates that RlmA, the enzyme responsible for methylating G 747, and RlmKL, the enzyme responsible for methylating G 2073, act before the accumulation of the 35S and 45S intermediates, results that agree with our previous Illumina NGS data ^10, 45^ ^34^. Furthermore, the data in this study reveal that stress conditions resulting from R331A DbpA expression do not modulate the levels of the m^1^G 747 and m^7^G 2073 modifications (Figure 2, Table S3, Table S4).

**Figure 2.**
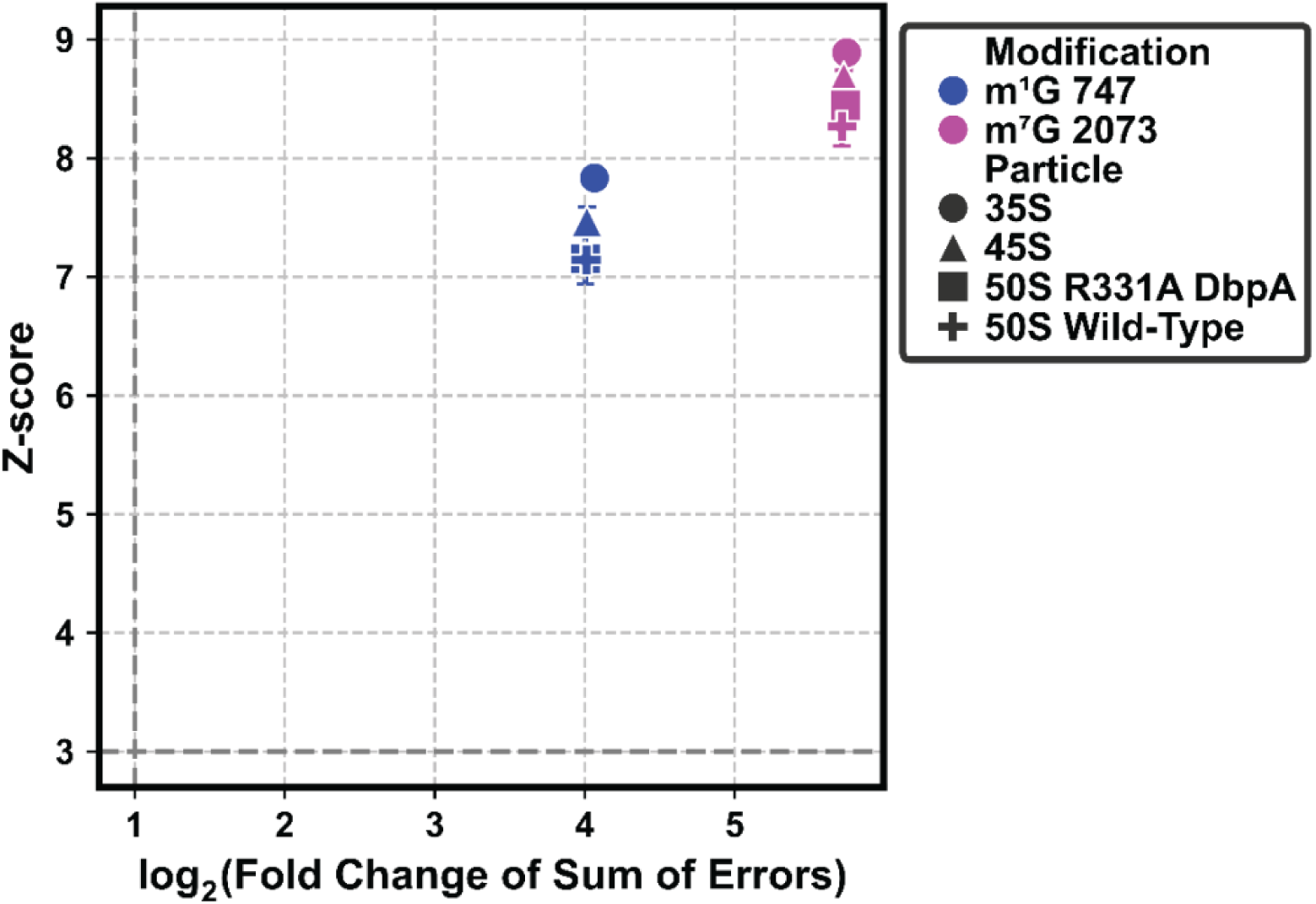
m^1^G and m^7^G modifications are incorporated to similar extents in 35S, 45S, and 50S ribosomal particles isolated from cells expressing either R331A or wild-type DbpA. Z-scores, Fold Changes, and errors shown in the plot were calculated as described in Figure 1. m^1^G modifications are shown in purple and m^7^G modifications in blue. Particle types are indicated as follows: 35S intermediates (circles), 45S intermediates (triangles), 50S LSU from cells expressing R331A DbpA (squares), and 50S LSU from cells expressing wild-type DbpA (crosses).

### R331A DbpA induced ribosome assembly stress does not affect incorporation of Ψ 748, Ψ 2461, and Ψ 2609

The Ψ 748 modification, located in helix 35 of 23S rRNA, is incorporated by the RluA enzyme^47^. Ψ 2461, located in the helix 89 of the PTC, is incorporated by the RluE enzyme and this modification is universally conserved ^48, 49^. Lastly, Ψ 2609 is incorporated by the RluB enzyme in helix 93 of the PTC ^48^. Helix 93 is a universally conserved structure implicated in peptide release ^50, 51^.

All of these modifications are present in the 35S and 45S intermediates at levels similar to those observed in the 50S subunits from cells expressing either R331A DbpA or wild-type DbpA (Figure 3, Table S2, Table S4). This reveals that these modifications are incorporated prior to the accumulation of the 35S and 45S intermediates in the cell. A conclusion which is in full agreement with our previous Illumina NGS data ^32, 34^.

**Figure 3.**
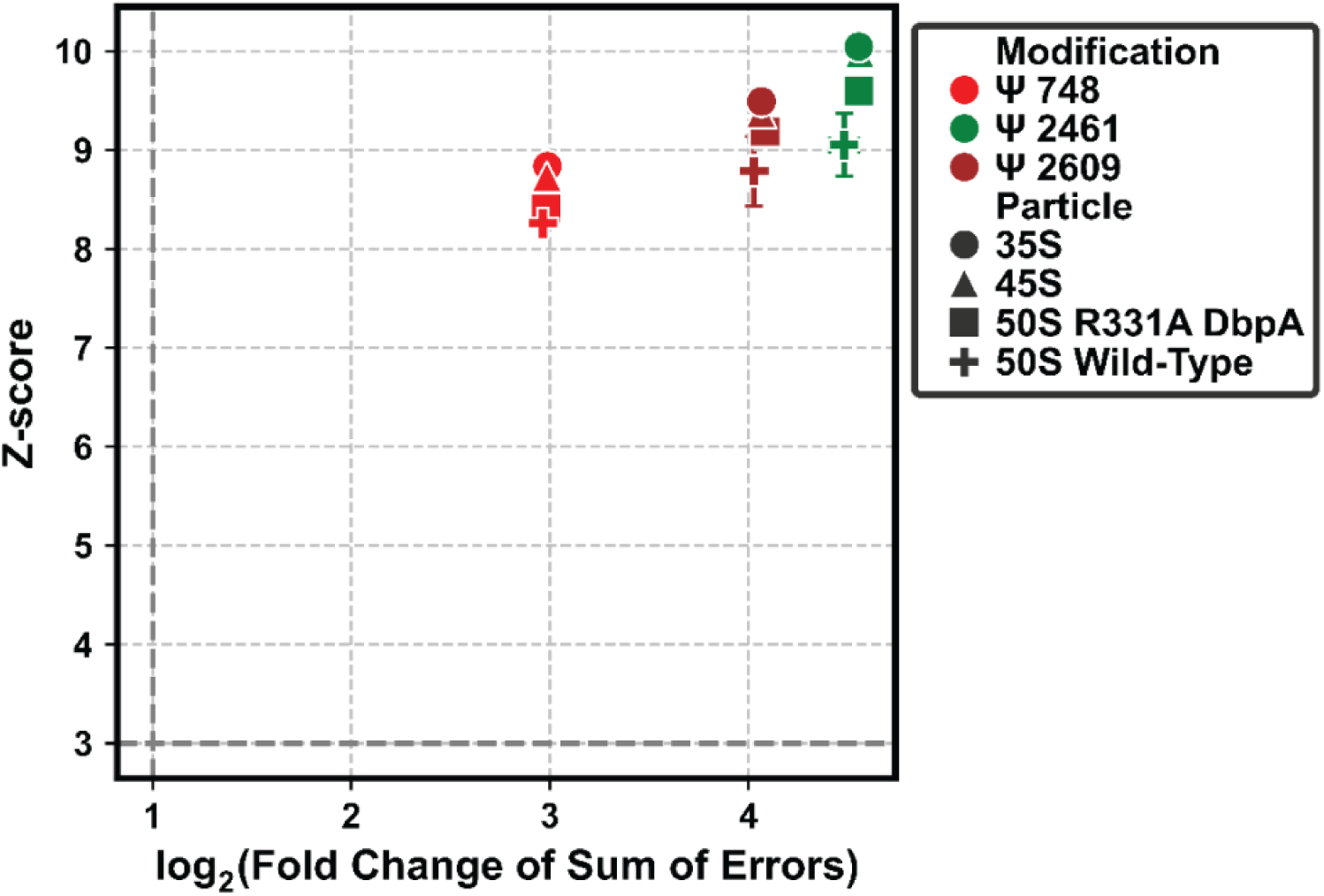
Ψ 748, Ψ 2461, Ψ 2609 have been incorporated to the same extent into the 35S, 45S, 50S from R331A expressing cells, and the 50S from the cells expressing wild-type DbpA. The Z-Scores, Fold Changes and errors shown in the plot are calculated as explained in Figure 1. Ψ 748 modifications are shown in red, and Ψ 2461 modifications in green, and Ψ 2609 maroon. Particle types are indicated as follows: 35S intermediates (circles), 45S intermediates (triangles), 50S LSU from cells expressing R331A DbpA (squares), and 50S LSU from cells expressing wild-type DbpA (crosses).

Furthermore, the levels of Ψ 748, Ψ 2461, Ψ 2609 modifications are also similar in the 50S subunits from cells expressing either R331A or wild type DbpA (Figure 3, Table S2, Table S3). In contrast, previous ONT DRS experiments revealed that Ψ 748 modification is decreased in *E. coli* cells subjected to cold shock, indicating that reprogramming of this modification is stress specific and occurs under certain environmental stresses but not under R331A DbpA induced ribosome assembly stress ^12^.

### The extent of late-incorporated helix 69 modifications, Ψ 1915, m^3^Ψ 1919, and Ψ 1921, is unchanged under R331A DbpA induced LSU assembly stress

The Ψ 1915, m^3^Ψ 1919, and Ψ 1921 modifications are located in helix 69, a universally conserved helix that forms intersubunit bridge B2a, and regulates ribosome stability, translation initiation, fidelity and translocation ^52–54^. Ψ 1915, m^3^Ψ 1919, and Ψ 1921 are incorporated by the RluD enzyme, and Ψ 1919 is subsequently methylated by the RlmH enzyme following formation of m^3^Ψ 1919 ^55, 56^. Two of these modifications, Ψ 1919 and Ψ 1921, are universally conserved ^55^. Deletion of *rluD*, which results in the absence of Ψ 1915, m^3^Ψ 1919, and Ψ 1921, leads to unstable 70S ribosomes, LSU assembly intermediates and translation termination errors ^55, 57–59^. Thus, helix 69 and the modifications it harbors are important for LSU biogenesis, stability, and translation accuracy ^52–54, 57, 58^. However, whether these modifications are reprogrammed under R331A DbpA induced stress within specific assembly pathways or in the mature 50S subunit remains to be determined. To address this, we examined the incorporation of helix 69 modifications in the 35S and 45S intermediates and the 50S LSU from cells expressing R331A DbpA or wild-type DbpA.

The Ψ 1915, m^3^Ψ 1919, and Ψ 1921 modifications are present in the 35S and 45S particles to a significantly lower extent than in the 50S LSU from the cells expressing wild-type DbpA (Figure 4, Table S2, Table S4). Hence, these modifications are not significantly incorporated in the 35S and 45S intermediates and are introduced during the later stages of LSU assembly. This conclusion is in complete agreement with our previous Illumina data collected in other LSU assembly intermediates ^14, 15, 32, 34, 60, 61^.

**Figure 4.**
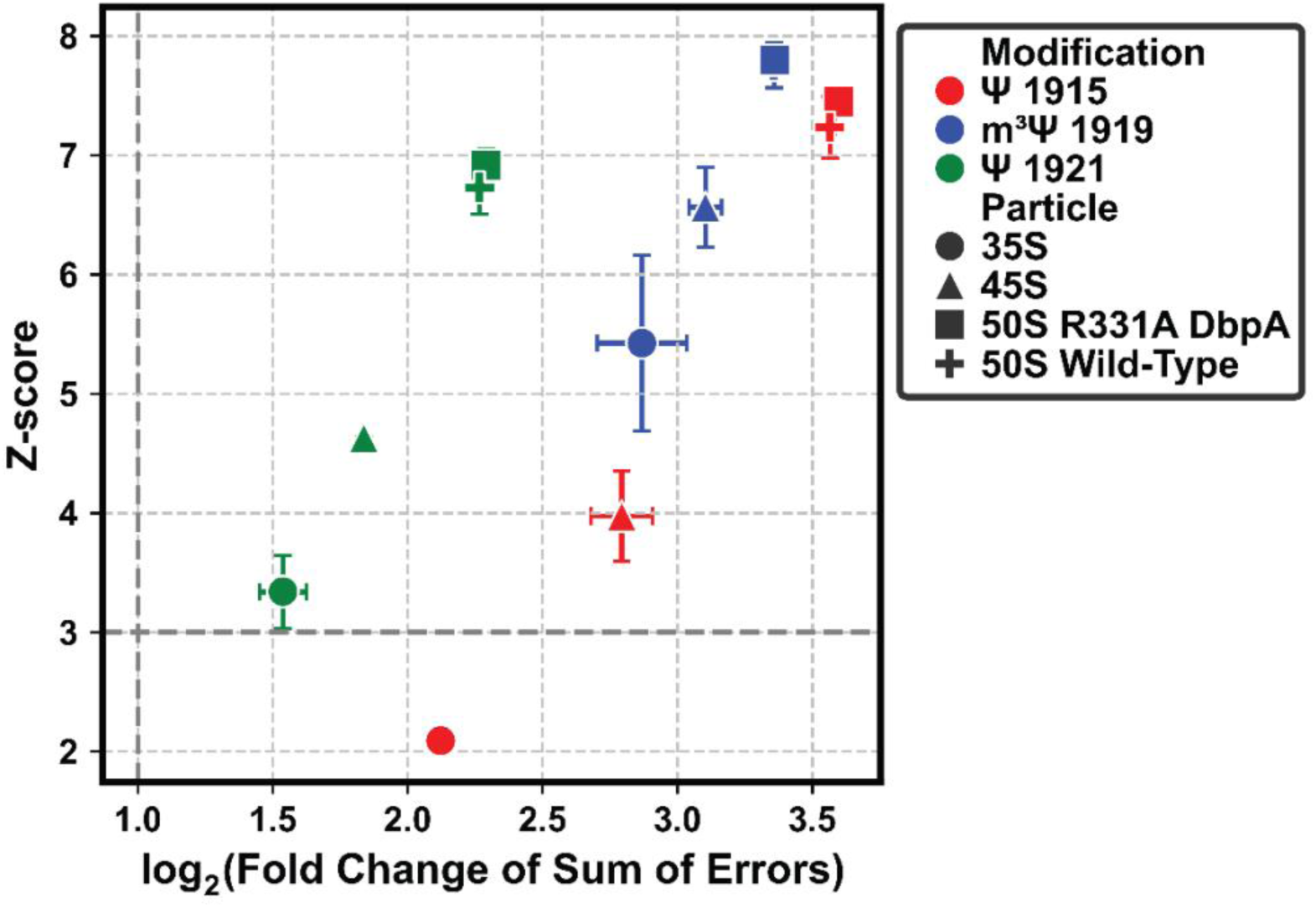
Late incorporation of Ψ 1915, m³Ψ 1919, and Ψ 1921 during LSU assembly is unaffected by assembly stress. Ψ 1915, m³Ψ 1919, and Ψ 1921 are incorporated in the 35S and 45S assembly intermediates to a significantly lower extent than in the mature 50S large subunit from cells expressing R331A DbpA. The extent of Ψ 1915, m^3^Ψ 1919, and Ψ 1921 modification in the 50S subunit is unaltered between cells expressing wild-type and R331A DbpA. Thus, R331A DbpA induced LSU assembly stress does not affect the final extent of Ψ 1915, m³Ψ 1919, and Ψ 1921 modification in the mature 50S LSU. Z-scores, fold changes, and associated errors shown in the plot were calculated as described in Figure 1. Ψ 1915 modifications are shown in red, and m^3^Ψ 1919 modifications in blue, and Ψ 1921 in green. Particle types are indicated as follows: 35S intermediates (circles), 45S intermediates (triangles), 50S LSU from cells expressing R331A DbpA (squares), and 50S LSU from cells expressing wild-type DbpA (crosses).

The extent of Ψ 1915, m^3^Ψ 1919, and Ψ 1921 modifications in the 50S subunit from wild-type and R331A DbpA expressing cells is similar (Figure 4, Table S2, Table S4). Thus, the ribosome assembly stress caused by R331A DbpA expression does not result in a change in the levels of Ψ 1915, m^3^Ψ 1919, and Ψ 1921 modifications. In contrast, the Ψ 1915, m^3^Ψ 1919, and Ψ 1921 modifications, as measured by ONT DRS, decrease in the 23S rRNA of cells exposed to cold shock, but not in cells subjected to metabolic stress ^12^. Collectively, our findings, and previous data, demonstrate that the final incorporation of Ψ 1915, m^3^Ψ 1919, and Ψ 1921 in the 50S is dynamically regulated in response to specific stress conditions ^12^.

Here, we determine only the extent of known RNA modifications. In the future, however, machine learning approaches could be employed to infer modification identity based on misincorporation, deletion, or insertion patterns observed in ONT DRS data, thereby distinguishing Ψ, m^3^Ψ, m^5^U, D and other uridine modifications. Our analysis of error patterns demonstrates that the m^3^Ψ modification predominantly produces deletion events in ONT DRS data, which is distinct from the error patterns associated with Ψ, m^5^U and D modifications in the SSU and LSU examined here (Figure S8). Determining m^3^Ψ error patterns across different sequence contexts may establish whether the ONT DRS signature of m^3^Ψ observed here can be generalized and employed to discriminate between different U modifications and m^3^Ψ in RNAs with unknown modification composition.

### Reduced m^2^A 2507 incorporation in the 50S under R331A DbpA induced assembly stress

The m^2^A 2507 modification, located in the PTC loop, is incorporated in *E. coli* by the RlmN enzyme, which is broadly conserved among bacteria ^31, 62–64^. Cells lacking RlmN exhibit reduced fitness, increased antibiotic sensitivity, and impaired translation accuracy ^63, 65^. In addition, recent studies have shown that the m^2^A 2507 modification increases the stability of 70S ribosomes and the lack of m^2^A 2507, in combination with Ψ 957, m^7^G 2073, m^2^G 2447, Ψ 2508, Ψ 2584, and Ψ 2601, also located within the PTC, affects translation elongation rate ^11, 66^. Collectively, these data indicate that the m^2^A 2507 modification regulates translation proofreading, elongation rate, ribosome maturation, and antibiotic resistance, providing a clear motivation for these experiments to determine how its incorporation is dynamically reprogrammed in response to physiological stress ^11, 63, 65, 66^.

The extent of m^2^A 2507 incorporation is lower in the 50S large subunit isolated from cells expressing the R331A DbpA mutant than in the 50S subunit from cells expressing wild-type DbpA (Figure 5, Table S3, Table S4). In addition, a recent ONT DRS study revealed that, in cells subjected to cold or metabolic stress, m^2^A 2507 incorporation into the 23S rRNA of the 50S subunit is reduced ^12^. Furthermore, during the stationary phase, characterized by nutrient limitation, the abundance of RlmN declines relative to that observed during exponential growth^67^. Together, these previous studies and our findings indicate that, under diverse stress conditions, the extent of m^2^A 2507 incorporation in *E. coli* is decreased. This reduction, which likely results in diminished translation rate, accuracy, and ribosome maturation, may be beneficial for cell survival under stress ^11, 65, 66^.

**Figure 5.**
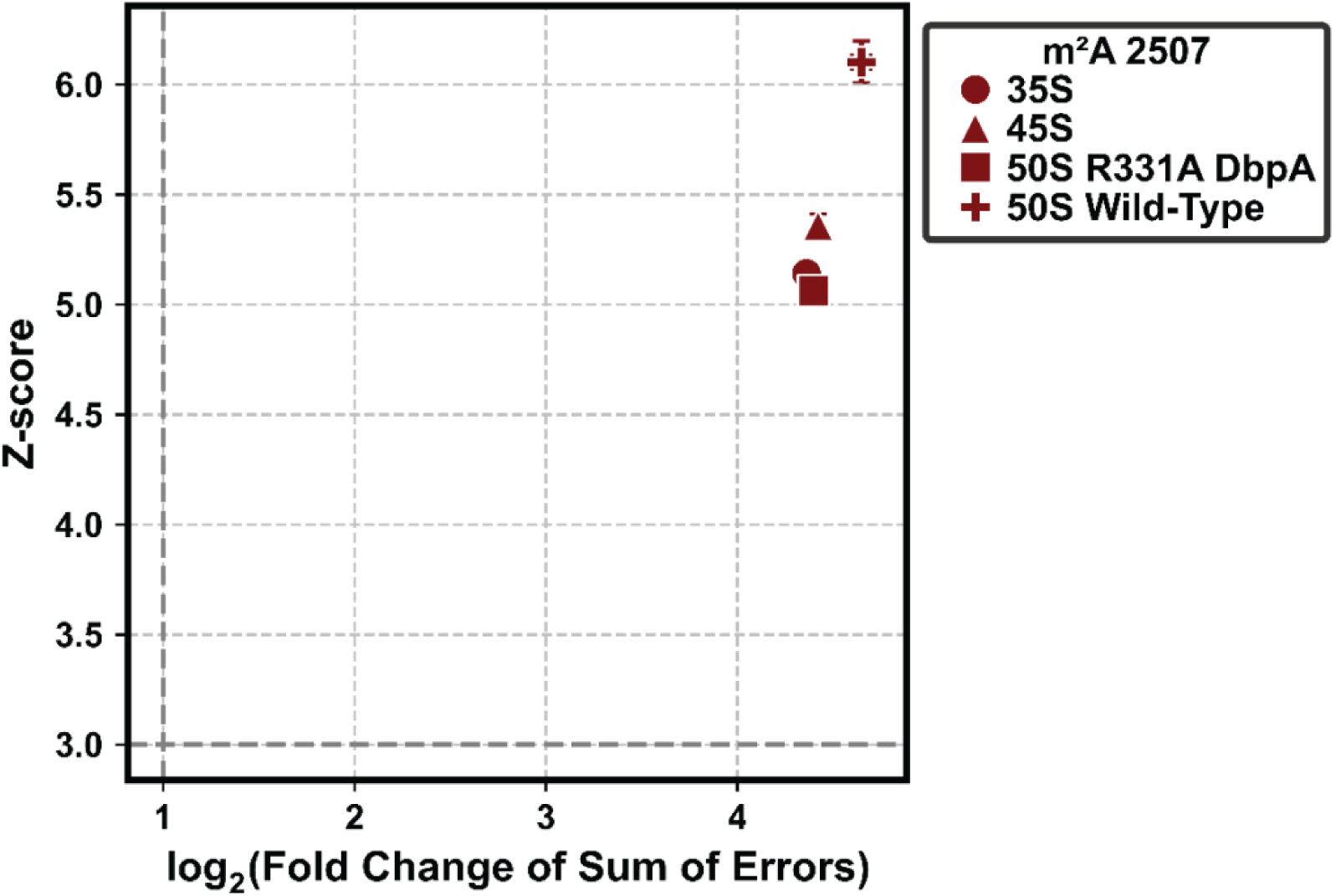
Decreased m²A 2507 incorporation in LSU assembly intermediates and the 50S subunit under R331A DbpA induced stress. Z-scores, fold-change values, and error rates were calculated as described in Figure 1. Particle types are indicated as follows: 35S intermediates (circles), 45S intermediates (triangles), 50S LSU from cells expressing R331A DbpA (squares), and 50S LSU from cells expressing wild-type DbpA (crosses).

### Differential incorporation of RluC dependent Ψ modifications under R331A DbpA induced LSU assembly stress

The RluC enzyme incorporates three Ψ modifications: Ψ 957, Ψ 2508, and Ψ 2584 ^68^. Ψ957 is located in helix 39, which forms intersubunit bridge B1a, Ψ 2508 is located in the peptidyl PTC loop, and Ψ 2584 is positioned in helix 90 of the PTC ^31, 46^. Thus, all three Ψ modifications are located in regions of the ribosome that are important for its stability and function. The presence of Ψ 957, Ψ 2508, and Ψ 2584 modifications promote LSU assembly ^11^. In addition, deletion of the *rluC* gene in a cell line that also lacks the *rlmE* methyltransferase gene results in a more severe decrease in translation elongation rate than deletion of *rlmE* alone ^66^. Lastly, the absence of Ψ 2508 renders cells sensitive to clindamycin, linezolid, and tiamulin, all of which target the PTC, where Ψ 2508 is located ^69^. Here, we assess how the Ψ 957, Ψ 2508, and Ψ 2584 modifications, which are important for ribosome stability, translation rate, and antibiotic resistance, are reprogrammed under ribosome assembly induced stress.

The levels of Ψ 957 and Ψ 2584 are similar in the 35S, 45S, and 50S particles from R331A DbpA expressing cells and in the 50S subunit from wild-type DbpA expressing cells. Thus, these modifications are incorporated by RluC prior to accumulation of 35S and 45S intermediates. A result that is in complete agreement with our previous Illumina NGS data (Figure 6, Table S2, Table S3) ^32^. Furthermore, R331A DbpA induced LSU assembly stress does not produce change in the final extent of Ψ 957 and Ψ 2584 incorporation into 50S LSU.

**Figure 6.**
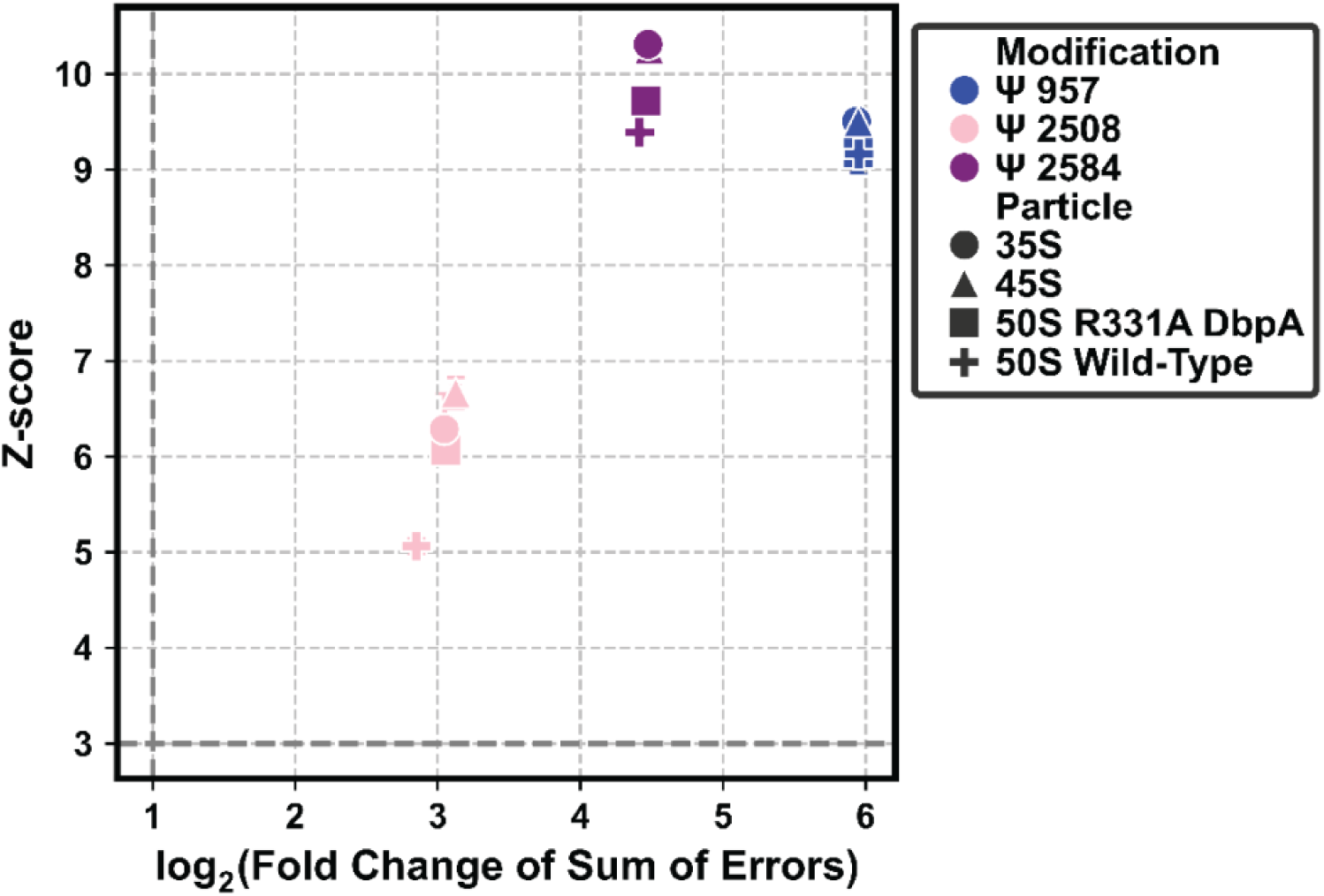
R331A DbpA induced ribosome assembly stress increases Ψ 2508 levels in the 50S LSU. The Ψ 2508 modification is incorporated to a greater extent in the 35S and 45S assembly intermediates and in mature 50S large subunits from cells expressing the R331A DbpA construct compared with the cells expressing the wild-type DbpA protein. The Ψ 2508 modification is incorporated by RluC enzyme, which also incorporates Ψ 957 and Ψ 2584. The levels of Ψ 957 and Ψ 2584 modifications are similar in the 50S LSU from cell expressing R331A or wild-type DbpA construct. Z-scores, fold changes, and associated errors were calculated as described in Figure 1. Ψ 957 modifications are shown in blue, Ψ 2508 modifications in pink and Ψ 2584 in purple. Particle types are indicated as follows: 35S intermediates (circles), 45S intermediates (triangles), 50S LSU from cells expressing R331A DbpA (squares), and 50S LSU from cells expressing wild-type DbpA (crosses).

In contrast, Ψ 2508 incorporation is increased in the 35S, 45S, and 50S particles from the R331A DbpA expressing cells compared with the 50S LSU from the cells expressing wild-type cells (Figure 6, Table S2, Tabel S4). First, these data demonstrate that Ψ 2508 modification is incorporated before 35S and 45S particle accumulation in R331A DbpA expressing cells, a conclusion that is in complete agreement with our previous Illumina NGS data ^32^. Second, the data presented here reveal that under ribosome assembly stress induced by R331A DbpA expression, the cell selectively increases the extent of the Ψ 2508 modification installed by RluC in 35S, 45S and 50S particles, while the levels of the other RluC dependent modifications, Ψ 957 and Ψ 2584, remain unchanged (Figure 6).

### RluF activity is reduced in distinct subsets of ribosome assembly pathways in cells expressing R331A DbpA

The Ψ 2608 modification is incorporated into the universally conserved helix 93 of PTC by the RluF enzyme ^48, 50^. Helix 93 is involved in peptide release ^51^. Ψ 2608 incorporation by RluF is diminished in the 50S LSU from cells expressing the R331A DbpA construct relative to 50S subunits from cells expressing wild-type DbpA (Figure 7, Table S2, and Table S4). Another ONT DRS study has shown that this modification decreases in 50S subunits from cells under cold-shock and increases under metabolic stress ^12^. Thus, the extent of Ψ2608 incorporation in this important PTC helix is dynamically regulated in response to specific stress conditions. This dynamic regulation, combined with the functional role of helix 93 in translation, suggests that Ψ 2608 may fine-tune ribosome function during stress adaptation by modulating structural elements involved in translation termination ^51^.

**Figure 7.**
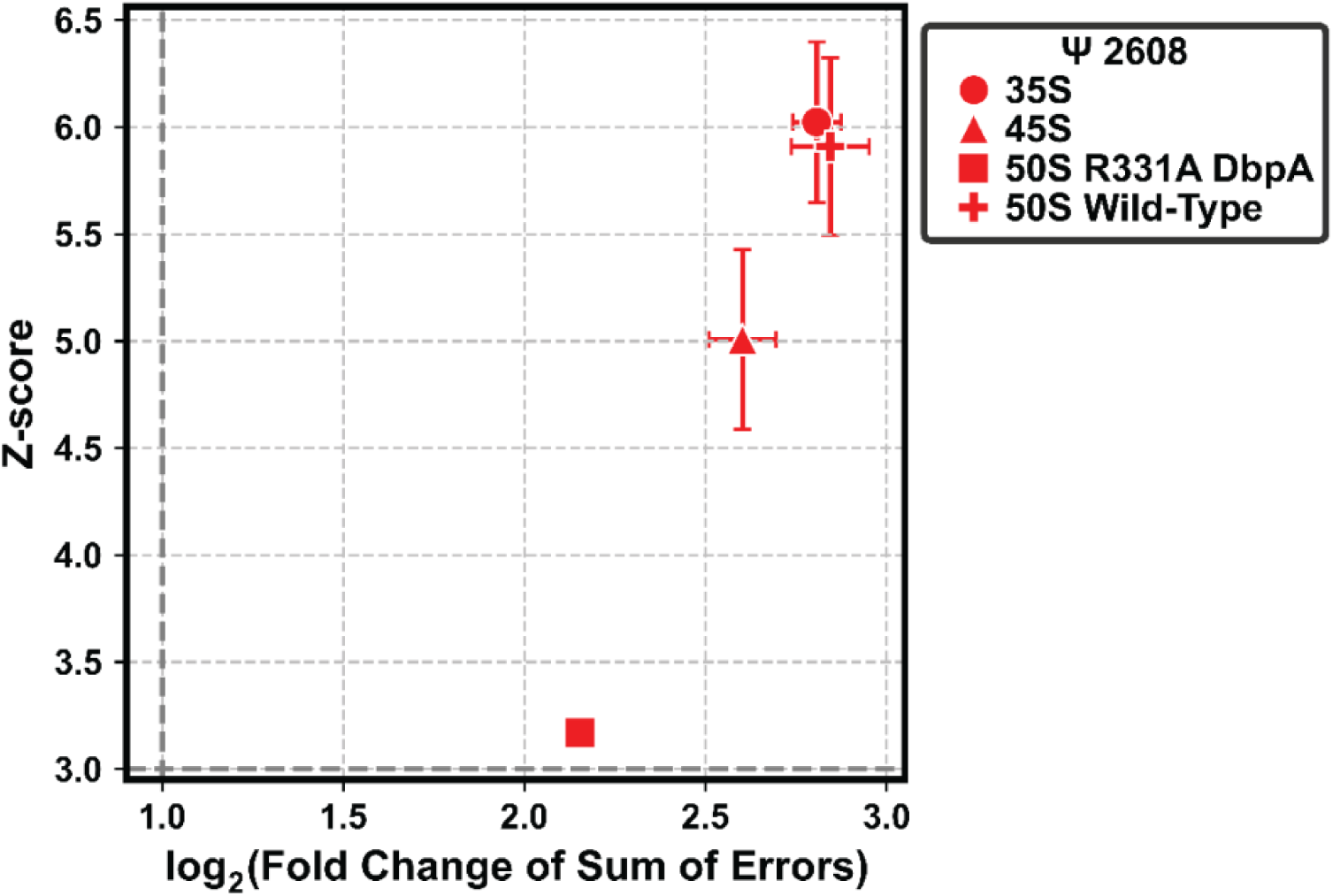
DbpA R331A induced assembly stress reduces Ψ2608 incorporation in distinct 50S LSU biogenesis pathways. The Ψ 2608 modification is incorporated at a lower extent in the 50S LSU isolated from the cells expressing the R331A DbpA construct compared with the 35S and 45S assembly intermediates and with the 50S LSU from the cells expressing wild-type DbpA. Because LSU biogenesis through the 35S and 45S intermediates represents only a subset of total LSU assembly pathways, these data indicate that under DbpA R331A induced ribosome assembly stress, Ψ 2608 incorporation is reduced in the LSU assembly pathways that do not involve 35S and 45S intermediate accumulation. Z-scores, fold changes, and associated errors were calculated as described in Figure 1. Particle types are indicated as follows: 35S intermediates (circles), 45S intermediates (triangles), 50S LSU from cells expressing R331A DbpA (squares), and 50S LSU from cells expressing wild-type DbpA (crosses).

Interestingly, Ψ 2608 is present at a lower level in the 50S LSU from R331A DbpA expressing cells than in the 35S and 45S intermediates from the same cells (Figure 7, Table S2, Table S4). Our previous *in vivo* rRNA pulse labeling studies determined that the pathways in which the 35S and 45S intermediates accumulate in cells expressing R331A DbpA account for approximately 40% of all ribosome assembly pathways ^28^. Thus, to produce 50S subunits with reduced Ψ 2608 modification, the cell may alter modification in at least a subset of the remaining 60% of assembly pathways. In other words, a fraction of the intermediates in these other pathways could become less favorable substrates for RluF.

The cells may favor this pathway specific regulation rather than reducing the overall abundance of RluF, because RluF not only modifies 23S rRNA at Ψ 2608 but also modifies tRNA^tyr^ at position 35 ^70^. By making a subset of ribosome assembly intermediates less accessible to RluF, the cell can selectively decrease Ψ 2608 modification in rRNA without interfering with its role in tRNA modification. In addition, this pathway specific regulation strategy enables switch-like regulation under stress conditions. When conditions change, the cell can rapidly increase 23S rRNA modification by RluF by restoring access to a larger fraction of assembly intermediates without a need to increase RluF production.

### The m^5^U 749 modification, introduced by RlmC, is incorporated at similar levels in 35S, 45S intermediates, and 50S from cells expressing wild type and R331A DbpA

The m^5^U 749 modification, which is incorporated by RlmC enzyme, is located in helix 35 ^71^. Helix 35 constitutes part of the peptide exit tunnel and the m^5^U 749 modification is highly conserved in bacteria ^46, 72^. We were unable to detect this modification in our previous Illumina NGS studies ^34, 35^. The data here show that this modification has been incorporated in the 35S and 45S intermediates to the same extent as in the 50S from cells expressing R331A (Figure 8A, Table S2, Table S4). Thus, this modification is incorporated before 35S and 45S intermediates are populated in cell. Furthermore, the extent of this modification has not been altered in the 50S large subunit of cells expressing R331A DbpA compared to the cells expressing wild-type DbpA (Figure 8A, Table S2, Table S4). Interestingly, a previous ONT DRS study observed the extent of m^5^U 749 modification decreased in *E. coli* cells exposed to cold shock stress but remains unchanged in cells exposed to metabolic stress ^12^. Therefore, m^5^U 749 is another modification which is dynamically regulated depending on the stress conditions.

**Figure 8.**
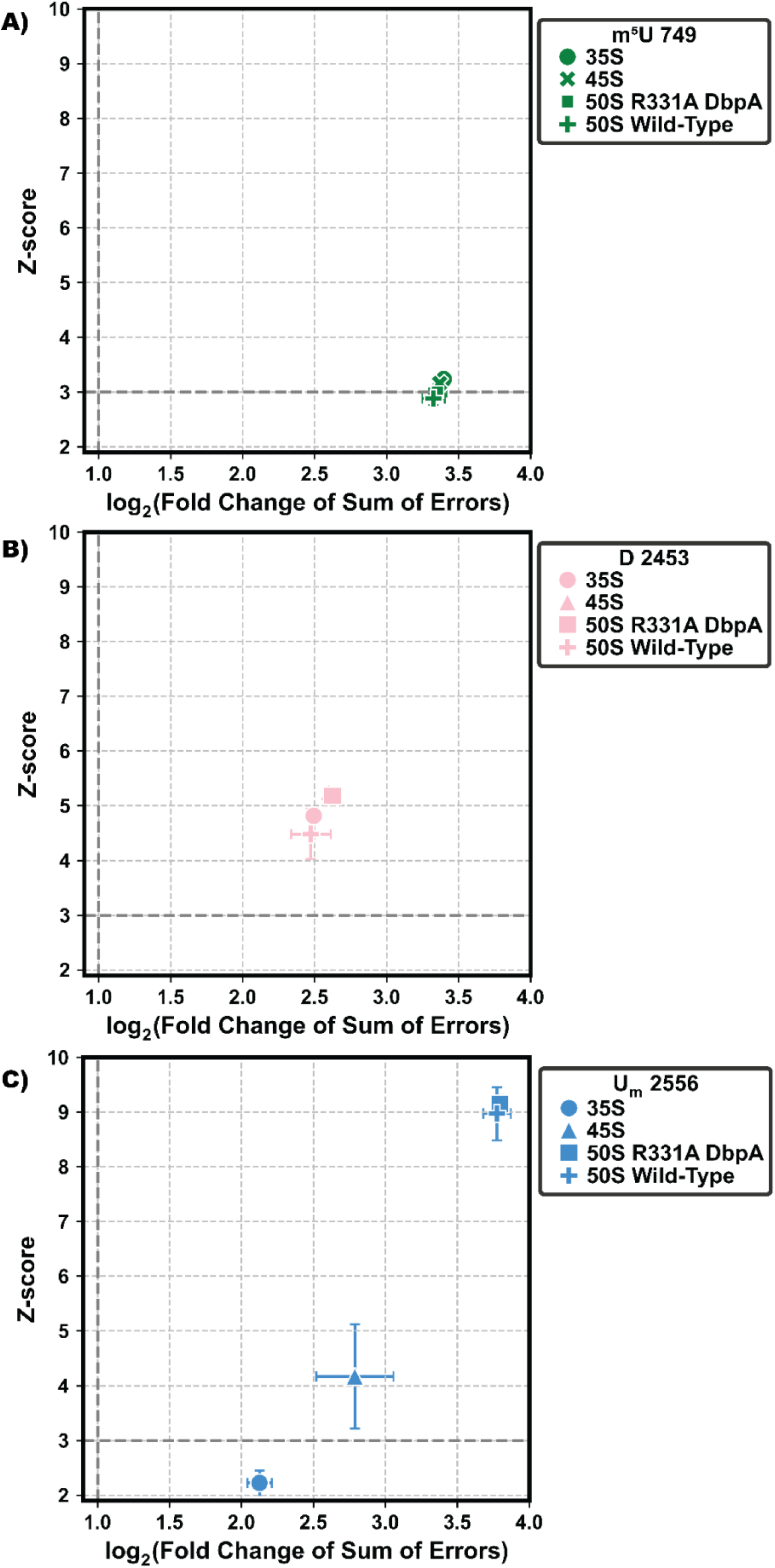
R331A DbpA induced ribosome biogenesis stress does not alter 50S rRNA modification levels of m^5^U 749, D 2453, or U_m_ 2556. A) m^5^U 749 is incorporated to a similar extent in 35S and 45S intermediates and in 50S subunits from cells expressing either wild-type or R331A DbpA. B) D 2453 is incorporated to a similar extent in the 35S and 45S intermediates and in 50S subunits cells expressing either wild-type or R331A DbpA. C) U_m_ 2556 is incorporated to a significantly lower extent in the 35S and 45S intermediates compared with 50S large subunits from cells expressing either R331A DbpA or wild-type DbpA. The levels of U_m_ 2556 are similar in the 50S isolated from cells expressing the wild-type or R331A DbpA. For panels A), B), and C), the Z-scores, fold changes, and associated errors shown in the plots were calculated as described in Figure 1. m^5^U 749 modifications are shown in green, D 2453 modifications in pink, U_m_ 2556 in blue. Particle types are indicated as follows: 35S intermediates (circles), 45S intermediates (triangles), 50S LSU from cells expressing R331A DbpA (squares), and 50S LSU from cells expressing wild-type DbpA (crosses).

### D 2453 is incorporated at similar extents in 50S particles from wild-type and R331A DbpA expressing cells

U 2453 is a universally conserved residue located in the PTC loop ^73^. Reduction of U 2453 to D 2453 is catalyzed by RdsA, an enzyme that is highly conserved across bacteria ^74^. A recent study further revealed that, in *E. coli* grown under anaerobic conditions, the 5′ carbon of D 2453 is methylated, which increases the stability of PTC and the efficiency of translation ^16^. Importantly, this change in methylation demonstrates that D 2453 is responsive to specific environmental stresses ^16^. Thus, we examined whether the extent of D 2453 modification is altered under R331A DbpA induced ribosome assembly stress.

The extent of D 2453 modification is similar in the 35S, 45S, and 50S particles from cells expressing R331A DbpA and in the 50S subunit from wild-type DbpA expressing cells (Figure 8B; Table S2; Table S4), indicating that the RdsA enzyme performs this modification prior to the accumulation of the 35S and 45S intermediates in the cell. This conclusion is in complete agreement with our previous Illumina NGS data ^34^.

Furthermore, the data presented here show that the extent of D 2453 modification in the 50S subunit is not altered under R331A DbpA induced ribosome assembly stress (Figure 8B, Table S2, Table S3). A previous ONT DRS study reported that the extent of D 2453 modification does not change under metabolic or cold-shock stress compared with cells grown at 37 °C in rich medium ^12^. Together, these findings indicate that D 2453 modification extent is largely insensitive to multiple ribosome assembly and environmental stress conditions, in contrast to anaerobic stress, under which 5′ methylation of D 2453 enhances PTC stability and translational efficiency ^12, 16^.

### The RlmE enzyme incorporates the U_m_ 2556 modification at similar levels in cells expressing either wild-type or R331A DbpA

The U_m_ 2556 modification, which is incorporated by RlmE enzyme in the A site of the ribosome, is highly conserved in all three kingdoms of life ^66, 75, 76^. Deletion of *rlmE* gene leads to the accumulation of a slowly rearranging 45S intermediate, indicating that U_m_ 2556 is directly involved in LSU assembly ^7^. Furthermore, 70S ribosomes lacking U_m_ 2556 exhibit a significantly reduced rate of elongation compared to ribosomes containing this modification ^66^. Its high conservation in all kingdoms of life, direct requirement for LSU maturation, and significant impact on translation elongation kinetics underscore U_m_ 2556 as a functionally important rRNA modification the regulation of which under diverse stress conditions merits further investigation ^7, 66, 75^.

Using Illumina NGS, we were unable to detect the U_m_ 2556 modification ^34^. In contrast, ONT DRS enabled robust detection of U_m_ 2556 in the 35S and 45S assembly intermediates, as well as in mature 50S large subunits isolated from cells expressing either wild-type or R331A DbpA (Figure 8C, Table S2, Table S4). The U_m_ 2556 levels in the 35S and 45S intermediates were significantly lower than those observed in the mature 50S subunits from both cell types. This comparison indicates that U_m_ 2556 installation by RlmE occurs predominantly at late stages of LSU assembly, after the accumulation of these intermediates in cell, consistent with previous reports ^14, 15, 29, 76^.

The overall extent of U_m_ 2556 modification in the 50S large subunit is unchanged in cells expressing the R331A DbpA construct compared to the cells expressing wild-type DbpA (Figure 8C, Table S2, S4). Similarly, U_m_ 2556 levels remain unchanged in *E. coli* exposed to metabolic or cold shock stress relative to cells grown in rich media at 37 °C ^12^. Although U_m_ 2556 levels could theoretically modulate ribosome assembly or translation, the profound ribosome biogenesis defects resulting from RlmE loss indicate that this modification may not be amenable to regulatory tuning ^7^. Accordingly, U_m_ 2556 levels remain invariant under all tested stress conditions (Figure 8C) ^12^.

### ONT DRS analysis of 3′-End rRNA processing in LSU assembly intermediates and the 50S from cells expressing wild-type or R331A DbpA

rRNA processing is an integral part of LSU maturation. Defects in this process can destabilize rRNA-protein interactions, compromise ribosome stability, and hinder the proper ribosome assembly, ultimately affecting cellular growth and protein synthesis efficiency ^77–79^. Here, we use ONT DRS to determine the 3’ end processing of 5S and 23S rRNA in the 35S, 45S and 50S particles from cells expressing R331A DbpA and in the 50S from cells expressing wild type DbpA.

In *E. coli*, the 5S, 16S and 23S rRNAs are transcribed as a single transcript, which is initially cleaved by RNase III into precursors of the 5S, 16S, and 23S rRNA molecules, each containing extra nucleotides at both the 5′ and 3′ ends ^41, 80–84^. After RNase III cleavage, RNase E trims both the 5′ and 3′ ends of the 5S rRNA precursor ^42, 85, 86^. At the 5′ end, the fragment resulting from RNase E cleavage may contain one, two, or three extra nucleotides, which are subsequently removed by RNase AM ^41^. At the 3′ end, the precursor may retain three or more extra nucleotides, which are trimmed by RNase T to produce the mature 3′ end ^42^.

In the case of 23S rRNA, the precursor formed by RNase III cleavage contains seven extra nucleotides at the 5′ end ^84^. The 5’ end is further trimmed to three nucleotides by an unidentified RNase and subsequently processed to the mature 5′ end by RNase AM ^41^. At the 3′ end, the 23S rRNA precursor contains eight extra nucleotides ^41, 84^. These are removed sequentially, first by RNase PH, RNase II, or other exonucleases, reducing the length to five extra nucleotides, and finally by RNase T, which generates the mature 3′ end of the 23S rRNA ^41^.

Our analysis of 5S and 23S rRNA is limited to the 3′ ends of these molecules because ONT DRS data analysis initiates at the 3′ end of RNA molecules and accurately reads 3′-terminal sequences but typically terminates approximately 10-15 nucleotides before the 5′ end is fully translocated through the pore ^87^. In addition, ONT DRS reads may end prematurely due to voltage fluctuations, pore unblocking events, or stalling of the motor protein that drives RNA translocation ^88–91^. As a result, ONT DRS is not suited for determining 5′-end processing.

Under our experimental conditions, we found that the 5S rRNA precursors generated by RNase E cleavage consistently contained five extra nucleotides at the 3′ end (Figure 9A, Figure S9). Notably, the 45S intermediate contained a larger fraction of incompletely processed 5S rRNA than the 35S intermediate. Moreover, incompletely processed 5S rRNA persisted in the 50S subunit from cells expressing R331A DbpA. Previous studies have shown that deletion of the *rnt* gene, which encodes RNase T and prevents complete 3′ end maturation of 5S rRNA, is not lethal to *E. coli* ^42^. Consistent with previous findings on *rnt* deletion, our results demonstrate that *E. coli* cells can survive with immature 3′ ends on 5S rRNA molecules ^42^.

**Figure 9.**
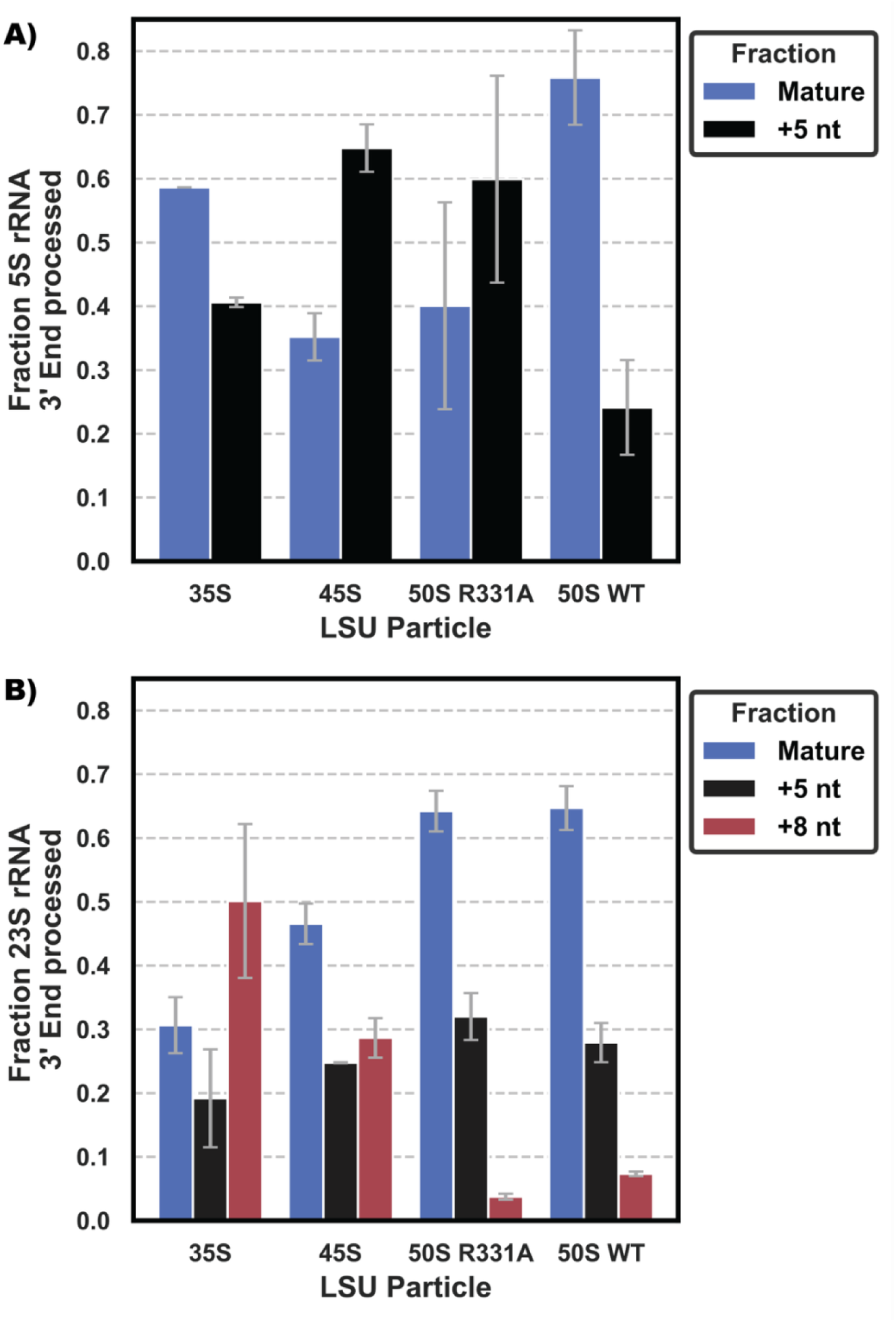
Differential 3′ end processing of 5S and 23S rRNA across ribosomal assembly intermediates and the 50S particles from cells expressing either wild-type (WT) or R331A DbpA. A) The 3′ end of 5S rRNA is more mature in the 35S intermediate than in the 45S intermediate. In 50S large subunits from R331A DbpA expressing cells, a substantial fraction of 5S rRNA molecules retain five additional nucleotides at their 3′ end, indicating incomplete processing. Blue bars depict the fraction of 5S rRNA molecules with correctly processed 3′ ends, red bars represent those with five extra nucleotides. B) In the R331A DbpA expressing cells, 23S rRNA in the 35S intermediate is less mature than in the 45S intermediate, and rRNA in the 45S intermediate is less mature than in the 50S LSU. Blue bars represent the fraction of 23S rRNA molecules with correctly processed 3′ ends. Black and red bars represent molecules containing five or eight unprocessed nucleotides at the 3′ end, respectively. The fractions of correctly processed and unprocessed 5S and 23S rRNAs were quantified as described in the Materials and Methods section.

Our data reveal that a larger fraction of 23S rRNA molecules are fully matured at the 3′ end in the 45S intermediate compared to the 35S intermediate, while the 23S rRNA in the 50S form the cells expressing wild-type DbpA and R331A are matured to a similar extent (Figure 9B, Figure S10). The 3′ end processing data here are in complete agreement with our previous 5′ end processing results, which showed that the 5′ end of 23S rRNA was also more extensively processed in the 45S intermediate than in the 35S intermediate ^28^. This consistency aligns with the established model that maturation of the 5′ and 3′ ends of 23S rRNA occurs concurrently^84^.

## CONCLUSION

Employing ONT DRS combined with EpiNano, we identified the following 23S rRNA modifications in the 35S and 45S intermediates, and in the 50S subunit, from cells expressing either wild-type or R331A DbpA constructs: m^1^G 747, Ψ 748, m^5^U 749, Ψ 957, Ψ 1915, m^3^Ψ 1919, Ψ 1921, m^7^G 2073, D 2453, Ψ 2461, m^2^A 2507, Ψ 2508, U_m_ 2556, Ψ 2584, Ψ 2608, and Ψ 2609 (Table S2, Table S4). Except for m^5^U 749 and U_m_ 2556, these modifications had been previously analyzed using Illumina NGS in the 35S and 45S intermediates and the 50S subunit from wild-type DbpA expressing cells ^32, 34^. This is the first study that analyzes the m^5^U 749 and U_m_ 2556 modifications in the 35S intermediate and m^5^U 749 modification in the 45S intermediate accumulating in cells expressing R331A DbpA (Figure 8A, Figure 8C) ^29, 32, 34^. Importantly, in here, with the objective of determining how R331A DbpA induced LSU assembly stress reprograms 23S rRNA modifications, we also assessed the above 16 modification levels in the 50S subunit from cells expressing the R331A DbpA protein and compared those levels to those in cells expressing wild-type DbpA.

Comparative analysis of 23S rRNA modification levels across LSU assembly intermediates and mature subunits revealed that m^2^A 2507 levels were decreased, while the Ψ 2508 were increased in cells expressing R331A DbpA when compared to the cells expressing wild type DbpA (Figure 5, Figure 6, Table S2, Table S4). In addition, Ψ 2608 was selectively downregulated in specific LSU assembly pathways in the R331A DbpA expressing cells (Figure 7, Table S2, Table S4). These findings demonstrate that expression of R331A DbpA, which induces LSU assembly stress, leads to the reprogramming of at least three rRNA modifications, either in a pathway-dependent manner during biogenesis or in the mature 50S subunit.

We propose that the selective access of modification enzymes within specific assembly pathways may serve as a regulatory mechanism by which cells modulate RNA modifications. Such a mechanism would allow dynamic control of RNA modification without altering enzyme abundance, enabling cells to rapidly restore modification patterns once homeostasis is reestablished.

Finally, we analyzed the 3′ end maturation of 5S and 23S rRNAs across the 35S and 45S intermediates and the 50S large subunit. Our data show that 23S rRNA is less processed in 35S intermediate when compared to the 45S, while 5S rRNA is more mature in the 35S intermediate than in the 45S (Figure 9). In the R331A DbpA expressing cells, the persistent immaturity of the 3′ end of 5S rRNA in the mature 50S subunit underscores how assembly associated stress can influence both the processing and modification landscapes of rRNA.

## DATA AVAILABILITY

The post-basecalling ONT DRS files, EpiNano and SAMTools processed files were deposited on Gene Expression Omnibus ^37, 39^. The accession code for these data is: GXXXXX

## ASSOCIATED CONTENT

### Supporting Information

Integrative Genomics Viewer (IGV) snapshots of alignments 23S rRNA sum of errors (SE) comparison of two biological replicates for the 35S (A), 45S (B), 50S isolated from the cells expressing R331A DbpA (C) and 50S isolated from the cells expressing wild-type (WT) DbpA (D. Comparison of 16S rRNA SE of two 30S replicates isolated from the cells expressing the wild-type (WT) DbpA protein. 23S rRNA ΔXᵢ comparison of two biological replicates for the 35S (A), 45S (B), 50S isolated from the cells expressing R331A DbpA (C) and 50S isolated from the cells expressing wild-type (WT) DbpA (D). 16S rRNA ΔXᵢ comparison of two 30S replicates isolated from the cells expressing wild-type (WT) DbpA. ΔXᵢ of each nucleotide in both biological replicates of 35S (A), 45S (B), 50S accumulated in cells expressing R331A DbpA (C), 50S accumulated in cells expressing wild-type (WT) DbpA (D). ΔXᵢ of each nucleotide in both biological replicates of 30S small subunit from the cells expressing wild-type (WT) DbpA protein. Basecalling error profiles for RNA modifications in the 30S and 50S particles accumulated in cells expressing wild-type DbpA. 3’ end read depth of mature and unprocessed 5S rRNA in 35S, 45S, 50S particles from R331A DbpA, and 50S from wild-type DbpA cells. 3’ end read depth of mature and unprocessed 23S rRNA in the 35S, 45S, 50S particles from R331A DbpA expressing cells, and the 50S from wild-type DbpA expressing cells. List of modifications and the enzymes that incorporate the modifications in 23S rRNA and 16S rRNA of E. coli. Z-Score and SE fold change for large subunit particles isolated from cells expressing either the wild-type DbpA or R331A construct. Z-Score and SE Fold Changes for the known modified positions present in the 16S rRNA of wild-type DbpA-expressing cells. 95% confidence interval (CI) of Z-scores and Fold Change for all detected modifications in 23S rRNA and 16S rRNA.

### Accession Code

*E. coli* DbpA Uniprot entry P21693; *E. coli* RluB Uniprot entry: P37765; *E. coli* RluC Uniprot entry: P0AA39; *E. coli* RluD Uniprot entry: P33643; *E. coli* RluF Uniprot entry: P32684; *E. coli* RlmH Uniprot entry: P0A818; *E. coli* RlmN Uniprot entry: P36979; *E. coli* RlmA Uniprot entry: P36999; *E. coli* RlmKL Uniprot entry:P75864; *E. coli* RdsA Uniprot entry: P3763; *E. coli* RlmE Uniprot entry P0C0R7; *E. coli* RlmC Uniprot P75817; *E. coli* DeaD P0A9P6.

## FUNDING

This work was supported in part by the National Institute of General Medical Sciences Grant R01- GM131062, the University of Texas System Rising STARs Program, and the start-up from the Chemistry and Biochemistry Department at the University of Texas at El Paso to E.K, and Beca de Posgrado en Ciencia y Humanidades al Extranjero from Secretaría de Ciencia, Humanidades, Tecnología e Innovación to L.A.G.M.

## ACKNOWLEDGMENTS

We are grateful to Dr. David Mohr from the Johns Hopkins University Genetic Core Facility, for help with the ONT DRS initial data analysis.

---

For Table of Contents use only

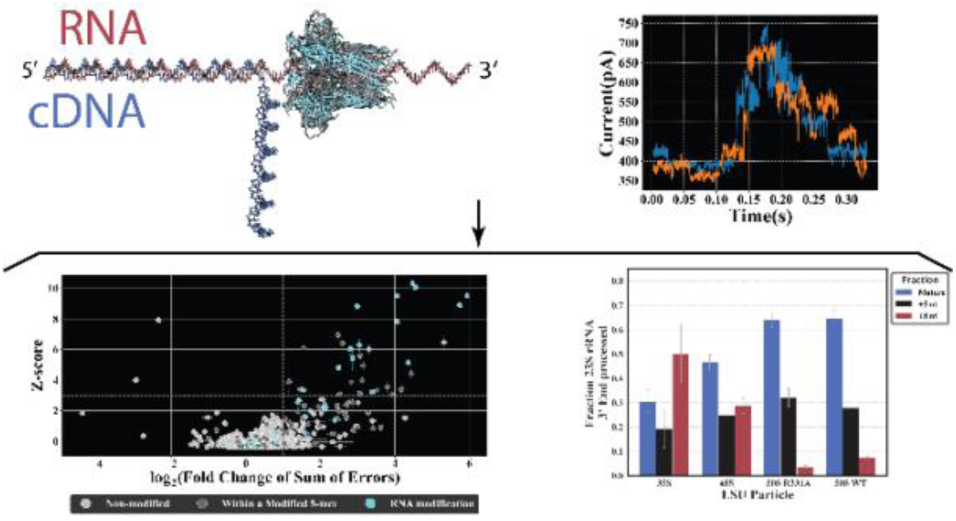

## Notes

### Competing Interest Statement

The authors have declared no competing interest.

## REFERENCES

[1] Toward Sequencing and Mapping of RNA Modifications Committee (Washington District of Columbia) (2024) Charting a future for sequencing rna and its modifications : a new era for biology and medicine, National Academies Press, Washington.

[2] Cui, L., Ma, R., Cai, J., Guo, C., Chen, Z., Yao, L., Wang, Y., Fan, R., Wang, X., and Shi, Y. (2022) RNA modifications: importance in immune cell biology and related diseases, Signal Transduct Target Ther 7, 334.

[3] Boo, S. H., and Kim, Y. K. (2020) The emerging role of RNA modifications in the regulation of mRNA stability, Exp Mol Med 52, 400–408.

[4] Gilbert, W. V., and Nachtergaele, S. (2023) mRNA Regulation by RNA Modifications, Annu Rev Biochem 92, 175–198.

[5] Chow, C. S., Lamichhane, T. N., and Mahto, S. K. (2007) Expanding the nucleotide repertoire of the ribosome with post-transcriptional modifications, ACS Chem Biol 2, 610–619.

[6] Boccaletto, P., and Baginski, B. (2021) MODOMICS: An Operational Guide to the Use of the RNA Modification Pathways Database, Methods Mol Biol 2284, 481–505.

[7] Arai, T., Ishiguro, K., Kimura, S., Sakaguchi, Y., Suzuki, T., and Suzuki, T. (2015) Single methylation of 23S rRNA triggers late steps of 50S ribosomal subunit assembly, Proc Natl Acad Sci U S A 112, E4707–4716.

[8] Gc, K., Gyawali, P., Balci, H., and Abeysirigunawardena, S. (2020) Ribosomal RNA Methyltransferase RsmC Moonlights as an RNA Chaperone, Chembiochem 21, 1885–1892.

[9] Keffer-Wilkes, L. C., Veerareddygari, G. R., and Kothe, U. (2016) RNA modification enzyme TruB is a tRNA chaperone, Proc Natl Acad Sci U S A 113, 14306–14311.

[10] Kimura, S., Ikeuchi, Y., Kitahara, K., Sakaguchi, Y., Suzuki, T., and Suzuki, T. (2012) Base methylations in the double-stranded RNA by a fused methyltransferase bearing unwinding activity, Nucleic Acids Res 40, 4071–4085.

[11] Ero, R., Leppik, M., Reier, K., Liiv, A., and Remme, J. (2024) Ribosomal RNA modification enzymes stimulate large ribosome subunit assembly in E. coli, Nucleic Acids Res 52, 6614–6628.

[12] Fleming, A. M., Bommisetti, P., Xiao, S., Bandarian, V., and Burrows, C. J. (2023) Direct Nanopore Sequencing for the 17 RNA Modification Types in 36 Locations in the E. coli Ribosome Enables Monitoring of Stress-Dependent Changes, ACS Chem Biol 18, 2211–2223.

[13] Delgado-Tejedor, A., Medina, R., Begik, O., Cozzuto, L., Lopez, J., Blanco, S., Ponomarenko, J., and Novoa, E. M. (2024) Native RNA nanopore sequencing reveals antibiotic-induced loss of rRNA modifications in the A- and P-sites, Nat Commun 15, 10054.

[14] Rabuck-Gibbons, J. N., Popova, A. M., Greene, E. M., Cervantes, C. F., Lyumkis, D., and Williamson, J. R. (2020) SrmB Rescues Trapped Ribosome Assembly Intermediates, J Mol Biol 432, 978–990.

[15] Siibak, T., and Remme, J. (2010) Subribosomal particle analysis reveals the stages of bacterial ribosome assembly at which rRNA nucleotides are modified, RNA 16, 2023–2032.

[16] Ishiguro, K., Midorikawa, K., Shigi, N., Kimura, S., Liiv, A., Yokoyama, T., Ito, T., Shirouzu, M., Remme, J., Miyauchi, K., and Suzuki, T. (2025) Hypoxia-induced ribosomal RNA modifications in the peptidyl-transferase center contribute to anaerobic growth of bacteria, Mol Cell.

[17] Fasnacht, M., Gallo, S., Sharma, P., Himmelstoss, M., Limbach, P. A., Willi, J., and Polacek, N. (2022) Dynamic 23S rRNA modification ho5C2501 benefits Escherichia coli under oxidative stress, Nucleic Acids Res 50, 473–489.

[18] Elles, L. M., and Uhlenbeck, O. C. (2008) Mutation of the arginine finger in the active site of Escherichia coli DbpA abolishes ATPase and helicase activity and confers a dominant slow growth phenotype, Nucleic Acids Res 36, 41–50.

[19] Tsu, C. A., and Uhlenbeck, O. C. (1998) Kinetic analysis of the RNA-dependent adenosinetriphosphatase activity of DbpA, an Escherichia coli DEAD protein specific for 23S ribosomal RNA, Biochemistry 37, 16989–16996.

[20] Diges, C. M., and Uhlenbeck, O. C. (2001) Escherichia coli DbpA is an RNA helicase that requires hairpin 92 of 23S rRNA, EMBO J 20, 5503–5512.

[21] Tsu, C. A., Kossen, K., and Uhlenbeck, O. C. (2001) The Escherichia coli DEAD protein DbpA recognizes a small RNA hairpin in 23S rRNA, RNA 7, 702–709.

[22] Karginov, F. V., and Uhlenbeck, O. C. (2004) Interaction of Escherichia coli DbpA with 23S rRNA in different functional states of the enzyme, Nucleic Acids Res 32, 3028–3032.

[23] Childs, J. J., Gentry, R. C., Moore, A. F., and Koculi, E. (2016) The DbpA catalytic core unwinds double-helix substrates by directly loading on them, RNA 22, 408–415.

[24] Moore, A. F., Gentry, R. C., and Koculi, E. (2017) DbpA is a region-specific RNA helicase, Biopolymers 107.

[25] Lopez de Victoria, A., Moore, A. F. T., Gittis, A. G., and Koculi, E. (2017) Kinetics and Thermodynamics of DbpA Protein’s C-Terminal Domain Interaction with RNA, ACS Omega 2, 8033–8038.

[26] Wurm, J. P., Glowacz, K. A., and Sprangers, R. (2021) Structural basis for the activation of the DEAD-box RNA helicase DbpA by the nascent ribosome, Proc Natl Acad Sci U S A 118.

[27] Ofengand, J., and Del Campo, M. (2004) Modified Nucleosides of Escherichia coli Ribosomal RNA, EcoSal Plus 1.

[28] Gentry, R. C., Childs, J. J., Gevorkyan, J., Gerasimova, Y. V., and Koculi, E. (2016) Time course of large ribosomal subunit assembly in E. coli cells overexpressing a helicase inactive DbpA protein, RNA 22, 1055–1064.

[29] Elles, L. M. S., Sykes, M. T., Williamson, J. R., and Uhlenbeck, O. C. (2009) A dominant negative mutant of the E. coli RNA helicase DbpA blocks assembly of the 50S ribosomal subunit, Nucleic Acids Res 37, 6503–6514.

[30] Shajani, Z., Sykes, M. T., and Williamson, J. R. (2011) Assembly of bacterial ribosomes, Annu Rev Biochem 80, 501–526.

[31] Ban, N., Nissen, P., Hansen, J., Moore, P. B., and Steitz, T. A. (2000) The complete atomic structure of the large ribosomal subunit at 2.4 A resolution, Science 289, 905–920.

[32] Koculi, E., and Cho, S. S. (2022) RNA Post-Transcriptional Modifications in Two Large Subunit Intermediates Populated in E. coli Cells Expressing Helicase Inactive R331A DbpA, Biochemistry 61, 833–842.

[33] Narayan, G., Gracia Mazuca, L. A., Cho, S. S., Mohl, J. E., and Koculi, E. (2023) RNA Post-transcriptional Modifications of an Early-Stage Large-Subunit Ribosomal Intermediate, Biochemistry 62, 2908–2915.

[34] Gracia Mazuca, L. A., Mohl, J. E., Cho, S. S., and Koculi, E. (2025) Post-Transcriptional Modifications of the Large Ribosome Subunit Assembly Intermediates in E. coli Expressing a Helicase-Inactive DbpA Variant, Biochemistry 64, 2976–2990.

[35] Koculi, E. G., R. C. (2025) High-throughput and single nucleotide resolution techniques for the determination of rna post-transcriptional modifications, In *US20190382832A1*, Assigned to UNIVERSITY OF CENTRAL FLORIDA RESEARCH FOUNDATION, INC, USA

[36] Li, H. (2018) Minimap2: pairwise alignment for nucleotide sequences, Bioinformatics 34, 3094–3100.

[37] Danecek, P., Bonfield, J. K., Liddle, J., Marshall, J., Ohan, V., Pollard, M. O., Whitwham, A., Keane, T., McCarthy, S. A., Davies, R. M., and Li, H. (2021) Twelve years of SAMtools and BCFtools, Gigascience 10.

38. [38] Institute, B. (2019) Picard Toolkit, Broad Institute, GitHub Repository.

[39] Liu, H., Begik, O., Lucas, M. C., Ramirez, J. M., Mason, C. E., Wiener, D., Schwartz, S., Mattick, J. S., Smith, M. A., and Novoa, E. M. (2019) Accurate detection of m(6)A RNA modifications in native RNA sequences, Nat Commun 10, 4079.

[40] Smola, M. J., Rice, G. M., Busan, S., Siegfried, N. A., and Weeks, K. M. (2015) Selective 2’-hydroxyl acylation analyzed by primer extension and mutational profiling (SHAPE-MaP) for direct, versatile and accurate RNA structure analysis, Nat Protoc 10, 1643–1669.

[41] Jain, C. (2020) RNase AM, a 5’ to 3’ exonuclease, matures the 5’ end of all three ribosomal RNAs in E. coli, Nucleic Acids Res 48, 5616–5623.

[42] Li, Z., and Deutscher, M. P. (1995) The tRNA processing enzyme RNase T is essential for maturation of 5S RNA, Proc Natl Acad Sci U S A 92, 6883–6886.

[43] Makhamreh, A., Tavakoli, S., Fallahi, A., Kang, X., Gamper, H., Nabizadehmashhadtoroghi, M., Jain, M., Hou, Y. M., Rouhanifard, S. H., and Wanunu, M. (2024) Nanopore signal deviations from pseudouridine modifications in RNA are sequence-specific: quantification requires dedicated synthetic controls, Sci Rep 14, 22457.

[44] Begik, O., Lucas, M. C., Pryszcz, L. P., Ramirez, J. M., Medina, R., Milenkovic, I., Cruciani, S., Liu, H., Vieira, H. G. S., Sas-Chen, A., Mattick, J. S., Schwartz, S., and Novoa, E. M. (2021) Quantitative profiling of pseudouridylation dynamics in native RNAs with nanopore sequencing, Nat Biotechnol 39, 1278–1291.

[45] Gustafsson, C., and Persson, B. C. (1998) Identification of the rrmA gene encoding the 23S rRNA m1G745 methyltransferase in Escherichia coli and characterization of an m1G745-deficient mutant, J Bacteriol 180, 359–365.

[46] Noeske, J., Wasserman, M. R., Terry, D. S., Altman, R. B., Blanchard, S. C., and Cate, J. H. (2015) High-resolution structure of the Escherichia coli ribosome, Nat Struct Mol Biol 22, 336–341.

[47] Wrzesinski, J., Nurse, K., Bakin, A., Lane, B. G., and Ofengand, J. (1995) A dual-specificity pseudouridine synthase: an Escherichia coli synthase purified and cloned on the basis of its specificity for psi 746 in 23S RNA is also specific for psi 32 in tRNA(phe), RNA 1, 437–448.

[48] Del Campo, M., Kaya, Y., and Ofengand, J. (2001) Identification and site of action of the remaining four putative pseudouridine synthases in Escherichia coli, RNA 7, 1603–1615.

[49] Tillault, A. S., Schultz, S. K., Wieden, H. J., and Kothe, U. (2018) Molecular Determinants for 23S rRNA Recognition and Modification by the E. coli Pseudouridine Synthase RluE, J Mol Biol 430, 1284–1294.

[50] Noller, H. F., Kop, J., Wheaton, V., Brosius, J., Gutell, R. R., Kopylov, A. M., Dohme, F., Herr, W., Stahl, D. A., Gupta, R., and Waese, C. R. (1981) Secondary structure model for 23S ribosomal RNA, Nucleic Acids Res 9, 6167–6189.

[51] Youngman, E. M., Brunelle, J. L., Kochaniak, A. B., and Green, R. (2004) The active site of the ribosome is composed of two layers of conserved nucleotides with distinct roles in peptide bond formation and peptide release, Cell 117, 589–599.

[52] Ali, I. K., Lancaster, L., Feinberg, J., Joseph, S., and Noller, H. F. (2006) Deletion of a conserved, central ribosomal intersubunit RNA bridge, Mol Cell 23, 865–874.

[53] Hirabayashi, N., Sato, N. S., and Suzuki, T. (2006) Conserved loop sequence of helix 69 in Escherichia coli 23 S rRNA is involved in A-site tRNA binding and translational fidelity, J Biol Chem 281, 17203–17211.

[54] Maivali, U., and Remme, J. (2004) Definition of bases in 23S rRNA essential for ribosomal subunit association, RNA 10, 600–604.

[55] Raychaudhuri, S., Conrad, J., Hall, B. G., and Ofengand, J. (1998) A pseudouridine synthase required for the formation of two universally conserved pseudouridines in ribosomal RNA is essential for normal growth of Escherichia coli, RNA 4, 1407–1417.

[56] Ero, R., Leppik, M., Liiv, A., and Remme, J. (2010) Specificity and kinetics of 23S rRNA modification enzymes RlmH and RluD, RNA 16, 2075–2084.

[57] Ejby, M., Sorensen, M. A., and Pedersen, S. (2007) Pseudouridylation of helix 69 of 23S rRNA is necessary for an effective translation termination, Proc Natl Acad Sci U S A 104, 19410–19415.

[58] Kipper, K., Sild, S., Hetenyi, C., Remme, J., and Liiv, A. (2011) Pseudouridylation of 23S rRNA helix 69 promotes peptide release by release factor RF2 but not by release factor RF1, Biochimie 93, 834–844.

[59] Gutgsell, N. S., Deutscher, M. P., and Ofengand, J. (2005) The pseudouridine synthase RluD is required for normal ribosome assembly and function in Escherichia coli, RNA 11, 1141–1152.

[60] Popova, A. M., and Williamson, J. R. (2014) Quantitative analysis of rRNA modifications using stable isotope labeling and mass spectrometry, J Am Chem Soc 136, 2058–2069.

[61] Nikolay, R., Hilal, T., Schmidt, S., Qin, B., Schwefel, D., Vieira-Vieira, C. H., Mielke, T., Burger, J., Loerke, J., Amikura, K., Flugel, T., Ueda, T., Selbach, M., Deuerling, E., and Spahn, C. M. T. (2021) Snapshots of native pre-50S ribosomes reveal a biogenesis factor network and evolutionary specialization, Mol Cell, 1200–1215.

[62] Kowalak, J. A., Bruenger, E., and McCloskey, J. A. (1995) Posttranscriptional modification of the central loop of domain V in Escherichia coli 23 S ribosomal RNA, J Biol Chem 270, 17758–17764.

[63] Toh, S. M., Xiong, L., Bae, T., and Mankin, A. S. (2008) The methyltransferase YfgB/RlmN is responsible for modification of adenosine 2503 in 23S rRNA, RNA 14, 98–106.

[64] Atkinson, G. C., Hansen, L. H., Tenson, T., Rasmussen, A., Kirpekar, F., and Vester, B. (2013) Distinction between the Cfr methyltransferase conferring antibiotic resistance and the housekeeping RlmN methyltransferase, Antimicrob Agents Chemother 57, 4019–4026.

[65] Benitez-Paez, A., Villarroya, M., and Armengod, M. E. (2012) The Escherichia coli RlmN methyltransferase is a dual-specificity enzyme that modifies both rRNA and tRNA and controls translational accuracy, RNA 18, 1783–1795.

[66] Bao, L., Liljeruhm, J., Crespo Blanco, R., Brandis, G., Remme, J., and Forster, A. C. (2024) Translational impacts of enzymes that modify ribosomal RNA around the peptidyl transferase centre, RNA Biol 21, 31–41.

[67] Terrazas-Lopez, M., Aitken, V., Zeczycki, T. N., and Koculi, E. (2025) Phase-Specific Antibiotic Resistance Mechanisms in an Escherichia coli B Strain, bioRxiv.

[68] Conrad, J., Sun, D., Englund, N., and Ofengand, J. (1998) The rluC gene of Escherichia coli codes for a pseudouridine synthase that is solely responsible for synthesis of pseudouridine at positions 955, 2504, and 2580 in 23 S ribosomal RNA, J Biol Chem 273, 18562–18566.

[69] Toh, S. M., and Mankin, A. S. (2008) An indigenous posttranscriptional modification in the ribosomal peptidyl transferase center confers resistance to an array of protein synthesis inhibitors, J Mol Biol 380, 593–597.

[70] Addepalli, B., and Limbach, P. A. (2016) Pseudouridine in the Anticodon of Escherichia coli tRNATyr(QPsiA) Is Catalyzed by the Dual Specificity Enzyme RluF, J Biol Chem 291, 22327–22337.

[71] Madsen, C. T., Mengel-Jorgensen, J., Kirpekar, F., and Douthwaite, S. (2003) Identifying the methyltransferases for m(5)U747 and m(5)U1939 in 23S rRNA using MALDI mass spectrometry, Nucleic Acids Res 31, 4738–4746.

[72] Auxilien, S., Rasmussen, A., Rose, S., Brochier-Armanet, C., Husson, C., Fourmy, D., Grosjean, H., and Douthwaite, S. (2011) Specificity shifts in the rRNA and tRNA nucleotide targets of archaeal and bacterial m5U methyltransferases, RNA 17, 45–53.

[73] O’Connor, M., Lee, W. M., Mankad, A., Squires, C. L., and Dahlberg, A. E. (2001) Mutagenesis of the peptidyltransferase center of 23S rRNA: the invariant U2449 is dispensable, Nucleic Acids Res 29, 710–715.

[74] Toubdji, S., Thullier, Q., Kilz, L. M., Marchand, V., Yuan, Y., Sudol, C., Goyenvalle, C., Jean-Jean, O., Rose, S., Douthwaite, S., Hardy, L., Baharoglu, Z., de Crecy-Lagard, V., Helm, M., Motorin, Y., Hamdane, D., and Bregeon, D. (2024) Exploring a unique class of flavoenzymes: Identification and biochemical characterization of ribosomal RNA dihydrouridine synthase, Proc Natl Acad Sci U S A 121, e2401981121.

[75] Nissley, A. J., Penev, P. I., Watson, Z. L., Banfield, J. F., and Cate, J. H. D. (2023) Rare ribosomal RNA sequences from archaea stabilize the bacterial ribosome, Nucleic Acids Res 51, 1880–1894.

[76] Caldas, T., Binet, E., Bouloc, P., Costa, A., Desgres, J., and Richarme, G. (2000) The FtsJ/RrmJ heat shock protein of Escherichia coli is a 23 S ribosomal RNA methyltransferase, J Biol Chem 275, 16414–16419.

[77] Gutgsell, N. S., and Jain, C. (2012) Gateway role for rRNA precursors in ribosome assembly, J Bacteriol 194, 6875–6882.

[78] Meskauskas, A., Baxter, J. L., Carr, E. A., Yasenchak, J., Gallagher, J. E., Baserga, S. J., and Dinman, J. D. (2003) Delayed rRNA processing results in significant ribosome biogenesis and functional defects, Mol Cell Biol 23, 1602–1613.

[79] Bennison, D. J., Irving, S. E., and Corrigan, R. M. (2019) The Impact of the Stringent Response on TRAFAC GTPases and Prokaryotic Ribosome Assembly, Cells 8.

[80] Dunn, J. J., and Studier, F. W. (1973) T7 early RNAs and Escherichia coli ribosomal RNAs are cut from large precursor RNAs in vivo by ribonuclease 3, Proc Natl Acad Sci U S A 70, 3296–3300.

[81] Young, R. A., and Steitz, J. A. (1978) Complementary sequences 1700 nucleotides apart form a ribonuclease III cleavage site in Escherichia coli ribosomal precursor RNA, Proc Natl Acad Sci U S A 75, 3593–3597.

[82] Bram, R. J., Young, R. A., and Steitz, J. A. (1980) The ribonuclease III site flanking 23S sequences in the 30S ribosomal precursor RNA of E. coli, Cell 19, 393–401.

[83] Nomura, M., Gourse, R., and Baughman, G. (1984) Regulation of the synthesis of ribosomes and ribosomal components, Annu Rev Biochem 53, 75–117.

[84] Gutgsell, N. S., and Jain, C. (2010) Coordinated regulation of 23S rRNA maturation in Escherichia coli, J Bacteriol 192, 1405–1409.

[85] Misra, T. K., and Apirion, D. (1979) RNase E, an RNA processing enzyme from Escherichia coli, J Biol Chem 254, 11154–11159.

[86] Roy, M. K., Singh, B., Ray, B. K., and Apirion, D. (1983) Maturation of 5-S rRNA: ribonuclease E cleavages and their dependence on precursor sequences, Eur J Biochem 131, 119–127.

[87] Ibrahim, F., Oppelt, J., Maragkakis, M., and Mourelatos, Z. (2021) TERA-Seq: true end-to-end sequencing of native RNA molecules for transcriptome characterization, Nucleic Acids Res 49, e115.

[88] Workman, R. E., Tang, A. D., Tang, P. S., Jain, M., Tyson, J. R., Razaghi, R., Zuzarte, P. C., Gilpatrick, T., Payne, A., Quick, J., Sadowski, N., Holmes, N., de Jesus, J. G., Jones, K. L., Soulette, C. M., Snutch, T. P., Loman, N., Paten, B., Loose, M., Simpson, J. T., Olsen, H. E., Brooks, A. N., Akeson, M., and Timp, W. (2019) Nanopore native RNA sequencing of a human poly(A) transcriptome, Nat Methods 16, 1297–1305.

[89] Soneson, C., Yao, Y., Bratus-Neuenschwander, A., Patrignani, A., Robinson, M. D., and Hussain, S. (2019) A comprehensive examination of Nanopore native RNA sequencing for characterization of complex transcriptomes, Nat Commun 10, 3359.

[90] Parker, M. T., Knop, K., Sherwood, A. V., Schurch, N. J., Mackinnon, K., Gould, P. D., Hall, A. J., Barton, G. J., and Simpson, G. G. (2020) Nanopore direct RNA sequencing maps the complexity of Arabidopsis mRNA processing and m(6)A modification, Elife 9.

[91] Jiang, F., Zhang, J., Liu, Q., Liu, X., Wang, H., He, J., and Kang, L. (2019) Long-read direct RNA sequencing by 5’-Cap capturing reveals the impact of Piwi on the widespread exonization of transposable elements in locusts, RNA Biol 16, 950–959.

