## SUPPORTING INFORMATION for "Direct RNA Sequencing Reveals Stress-Dependent and Pathway-Specific rRNA Modification Reprogramming During 50S Biogenesis"

______________________

Isaac S. Weislow presently is affiliated with Department of Chemistry, University of Washington, Seattle, Washington 98195, United States of America

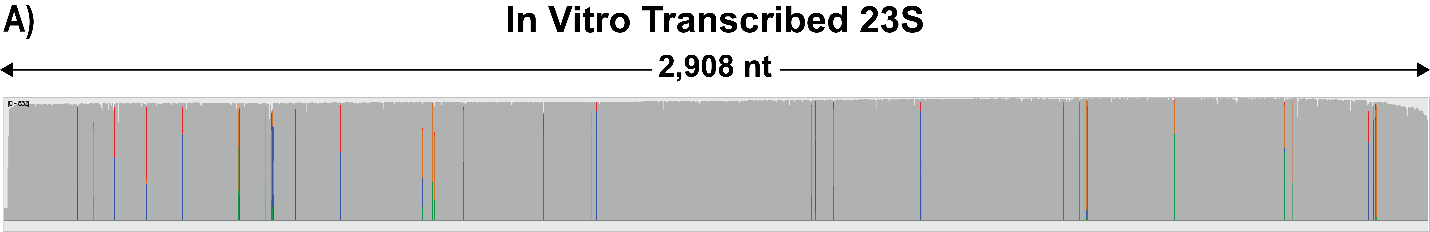

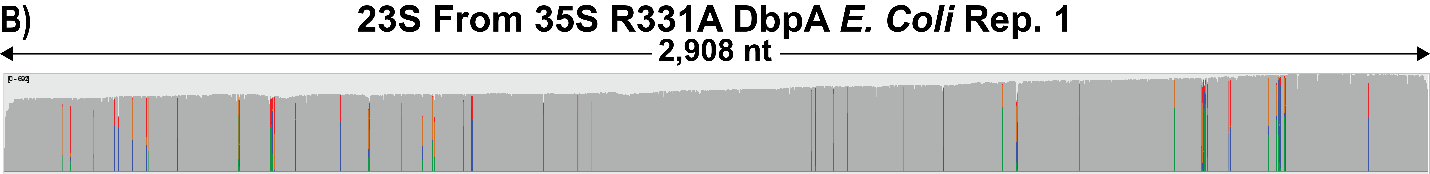

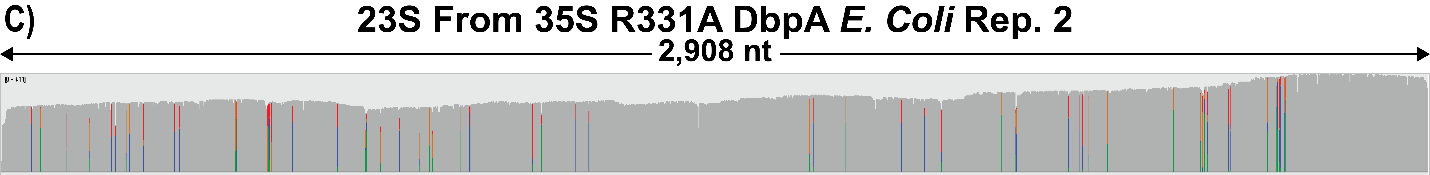

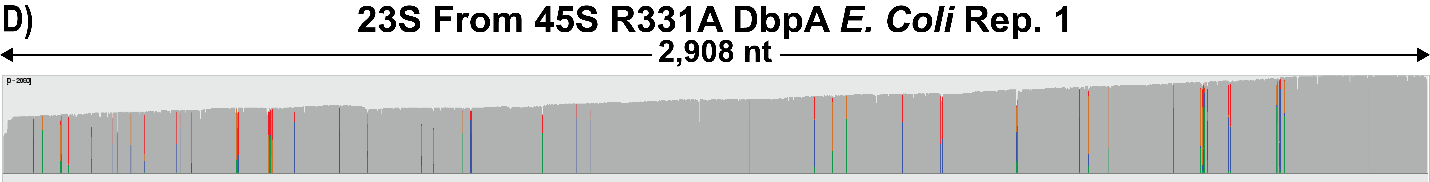

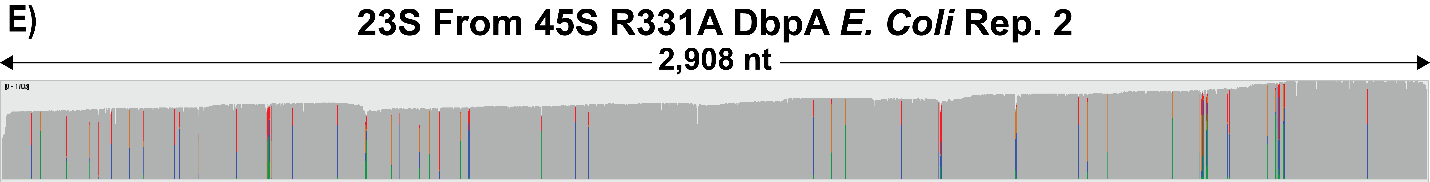

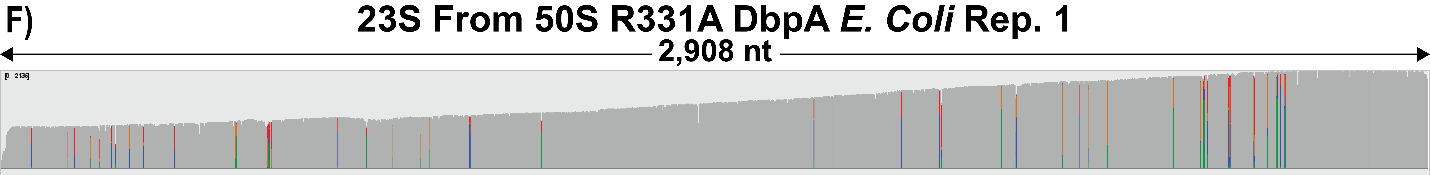

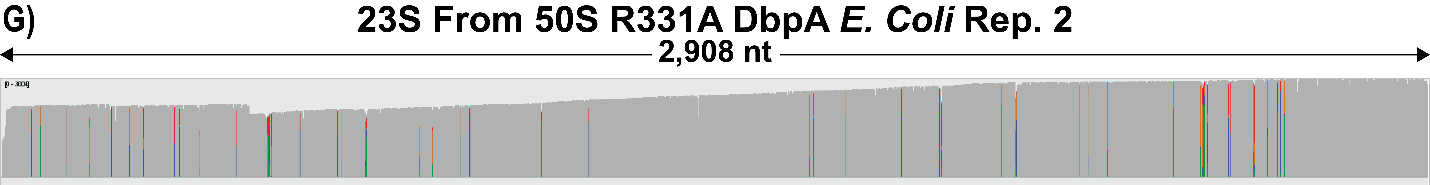

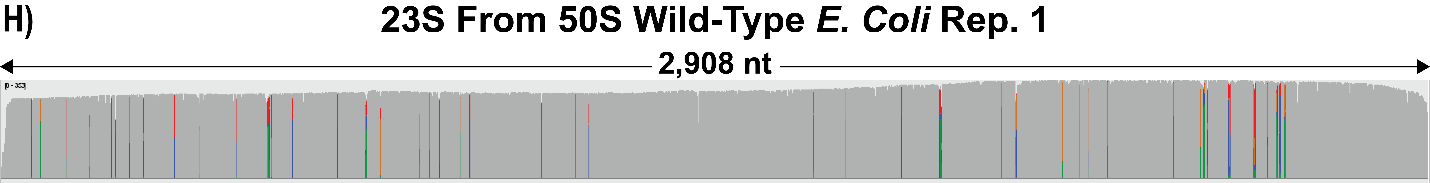

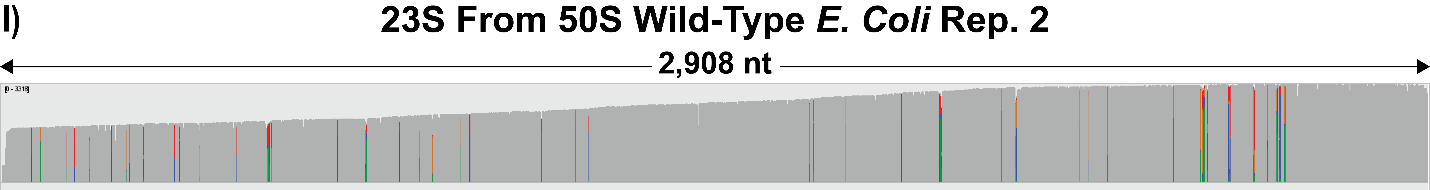

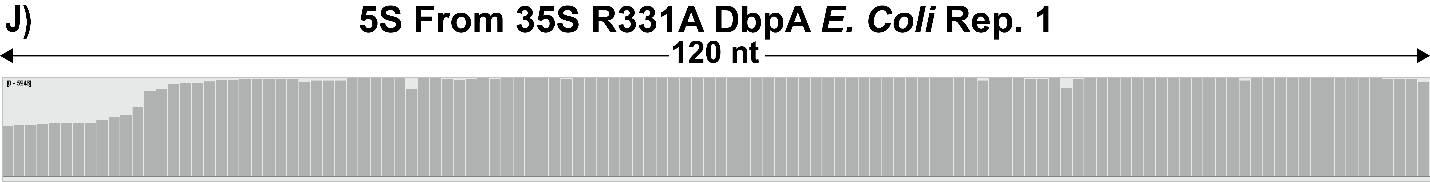

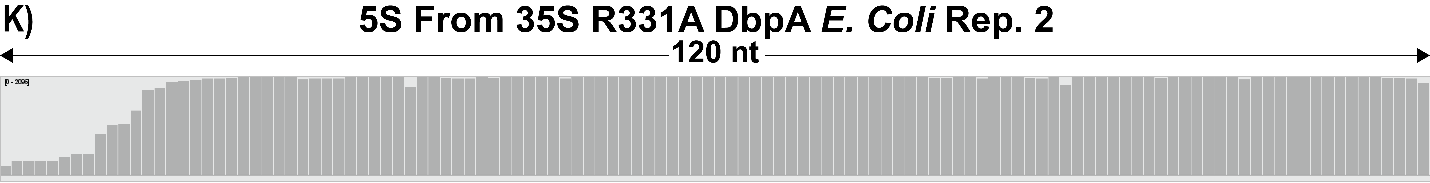

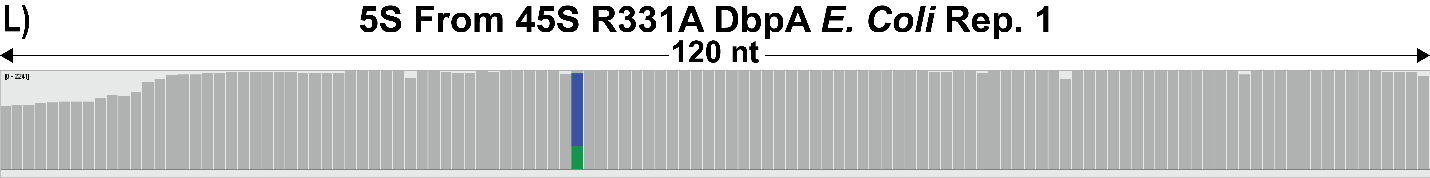

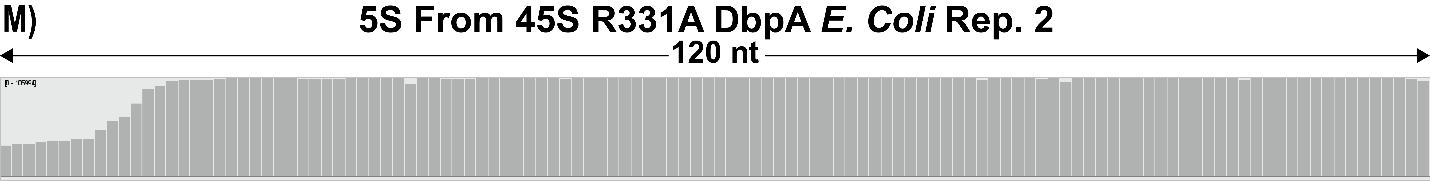

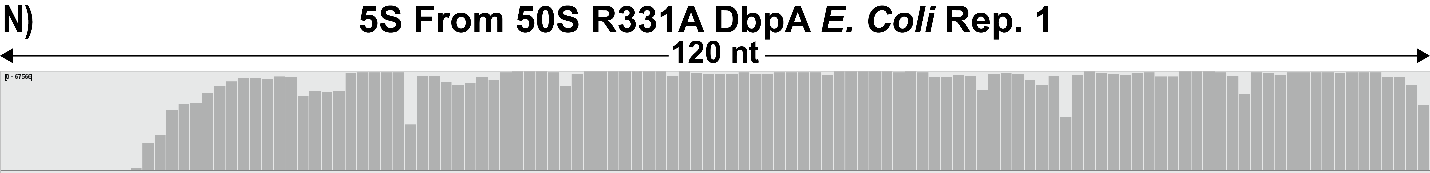

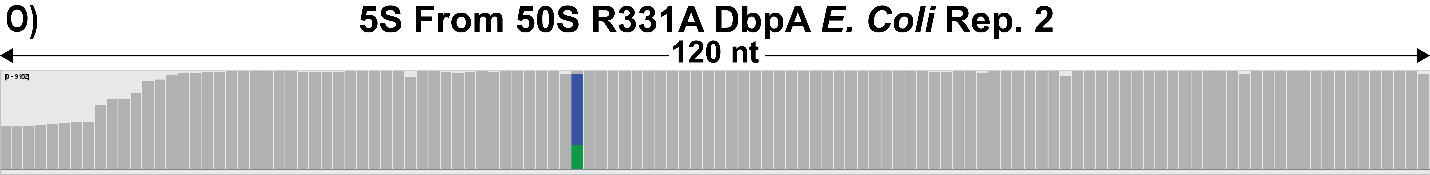

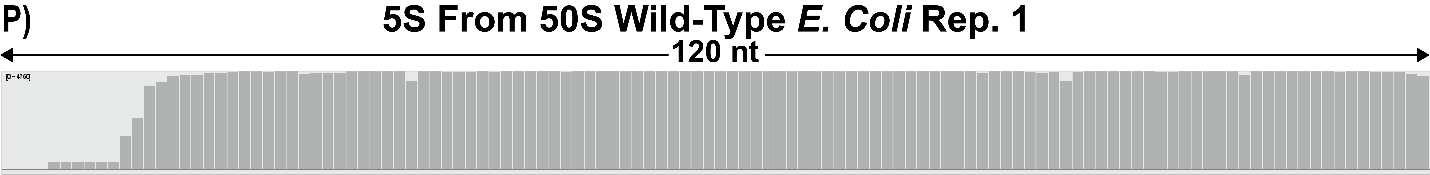

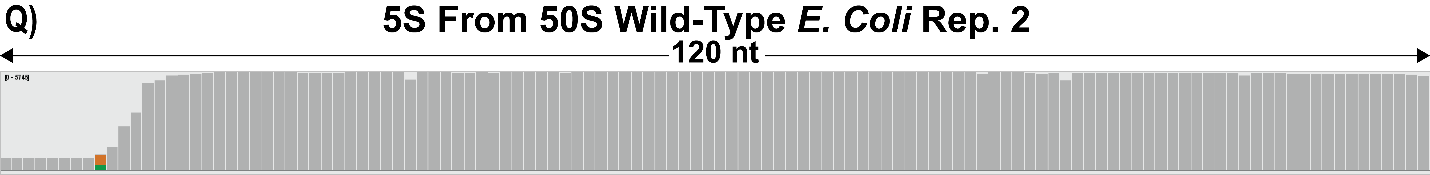

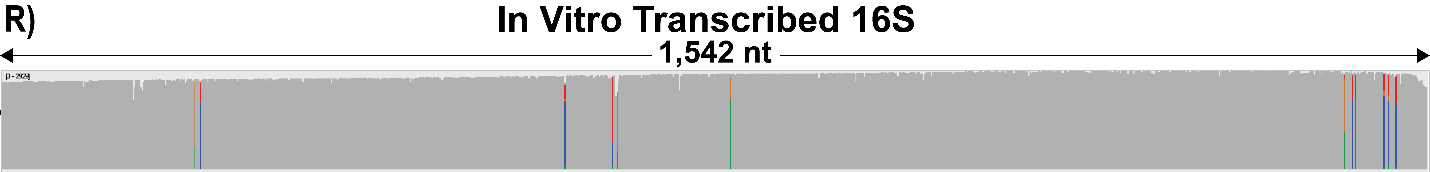

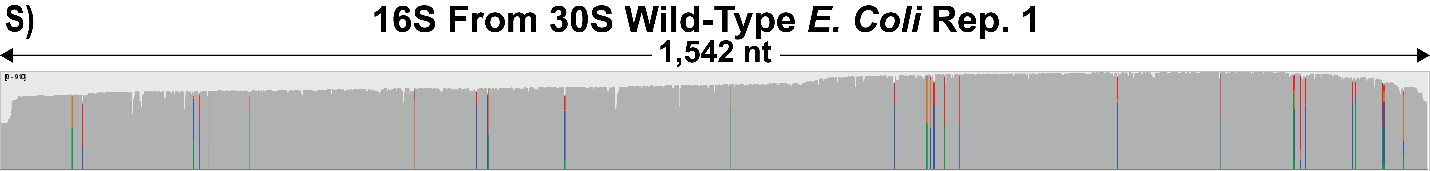

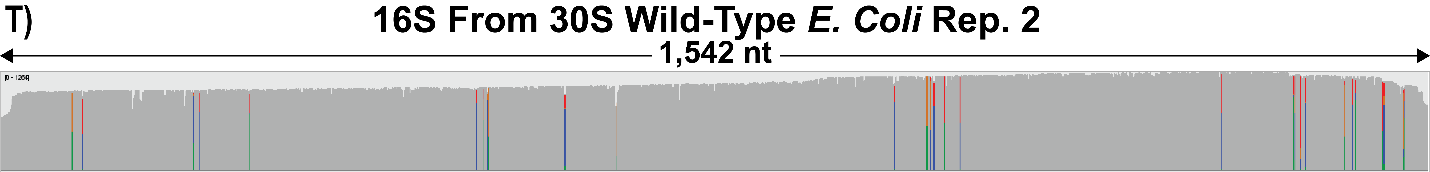

**Figure S1. Integrative Genomics Viewer (IGV) snapshots of alignments***^1^***.** Panels (A) to (T) depict the alignments performed with Minimap2 for each of the different RNA samples analyzed in this study *^2^*. Each alignment is labeled with the name of the ribosomal particle the 23S rRNA or 16S rRNA were isolated from and the biological sample to which it corresponds (Rep. 1 for biological replicate 1, and Rep. 2 for biological replicate 2). The *in-vitro* transcribed 16S and 23S rRNA were produced from DNA plasmids as detailed in the manuscript. The sequences were depicted in log scale. Gray bars represent positions where the base called was a match to the reference sequence. Color bars represent positions where the base called had a mismatch frequency greater than 0.2 *^1^*. The color scheme for the mismatches with a frequency greater than 0.2 is as follows: A green portion of the bar represents a mismatch to an A; a red portion of the bar represents a mismatch to a U; a blue portion of the bar represents a mismatch to a C; and an orange portion of the bar represents a mismatch to a G*^1^*.

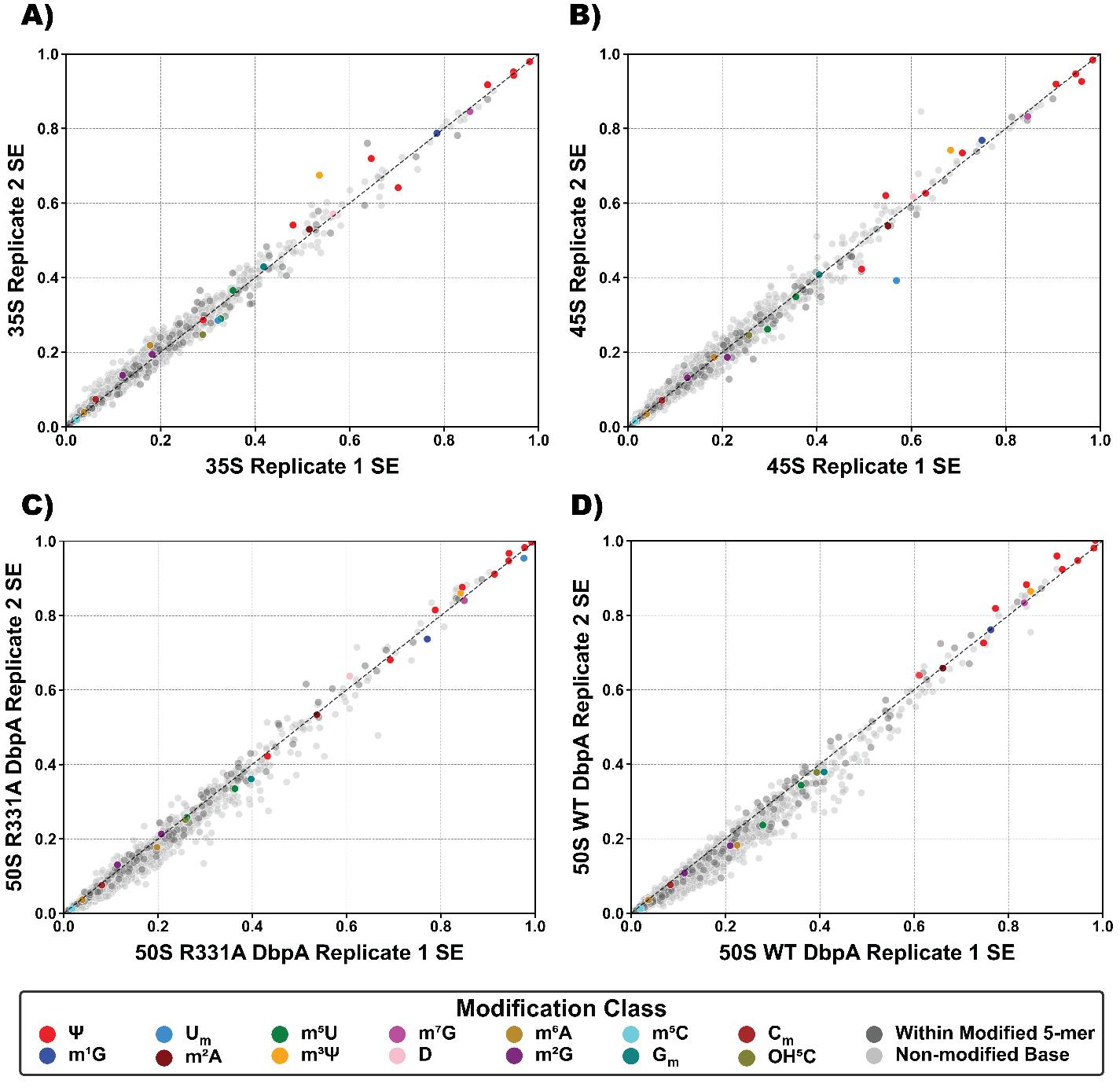

**Figure S2. 23S rRNA sum of errors (SE) comparison of two biological replicates for the 35S (A), 45S (B), 50S isolated from the cells expressing R331A DbpA (C) and 50S isolated from the cells expressing wild-type (WT) DbpA (D**. The SE is obtained from EpiNano as explained in the Materials and Methods section of the manuscript *^3^*. The nucleotides with no known modifications are shown in silver color, the nucleotides within five nucleotides of a known modification are shown as gray, the Ψ are shown as red, m^1^G as dark blue, U_m_ as sky blue, m^2^A as maroon, m^5^U as green, m^3^Ψ as orange, m^7^G as magenta, D as pink, m^6^A as golden, m^2^G as purple, m^5^C as cyan, G_m_ as teal, C_m_ as dark red, and OH^5^C as olive.

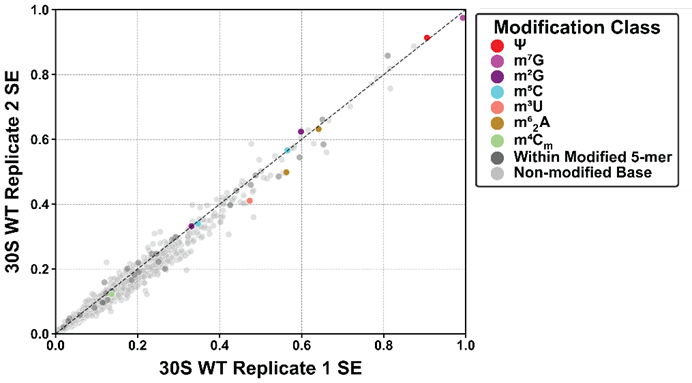

**Figure S3. Comparison of 16S rRNA SE of two 30S replicates isolated from the cells expressing the wild-type (WT) DbpA protein.** The SE is obtained from EpiNano as explained in the Materials and Methods section of the manuscript *^3^*. The nucleotides with no known modifications are shown in silver color, the nucleotides within five nucleotides of a known modification are shown in gray, Ψ nucleotides are shown as red, m^7^G in magenta, m^2^G in dark purple, m^5^C in cyan, m^3^U in salmon pink, m^6^_2_A in gold, and m^4^C_m_ in light green.

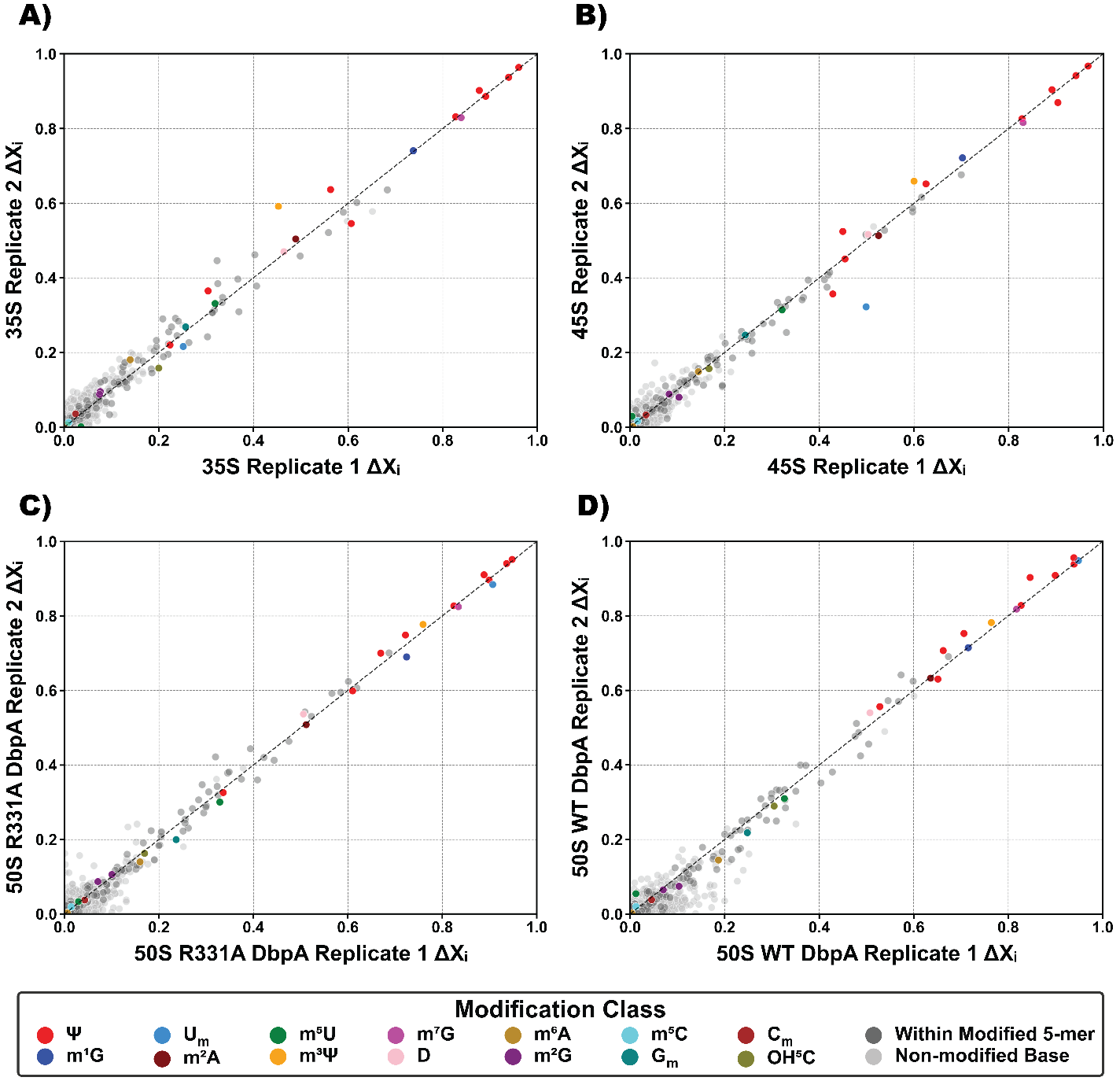

**Figure S4. 23S rRNA ΔXᵢ comparison of two biological replicates for the 35S (A), 45S (B), 50S isolated from the cells expressing R331A DbpA (C) and 50S isolated from the cells expressing wild-type (WT) DbpA (D).** ΔXᵢ was calculated as explained in the Materials and Methods section of the paper. The nucleotides with no known modifications are shown in silver color, the nucleotides within five nucleotides of a known modification are shown as gray, Ψ are shown as red, m^1^G as dark blue, U_m_ as sky blue, m^2^A as maroon, m^5^U as green, m^3^Ψ as orange, m^7^G as magenta, D as pink, m^6^A as golden, m^2^G as purple, m^5^C as cyan, G_m_ as teal, C_m_ as dark red, and OH^5^C as olive.

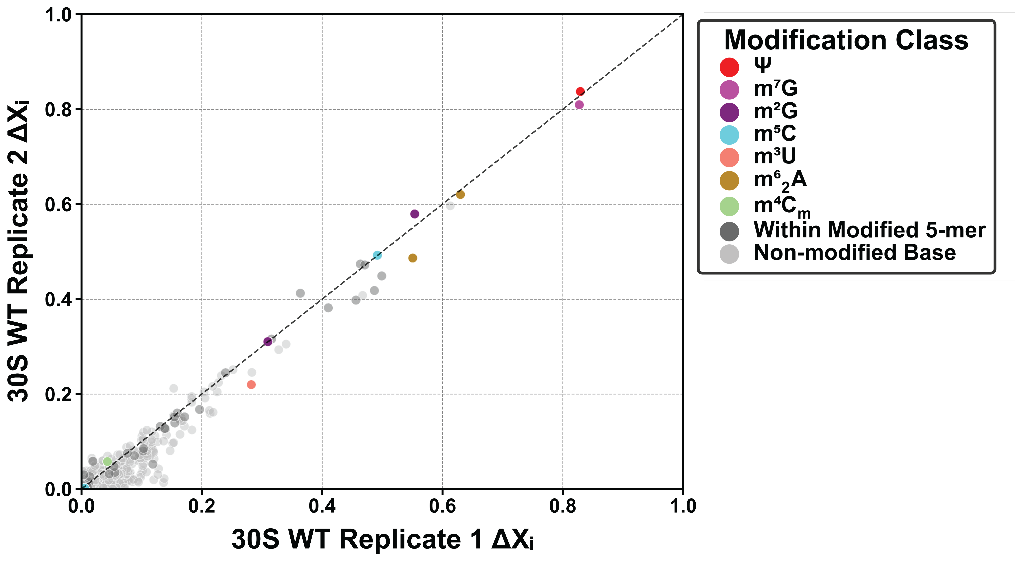

**Figure S5. 16S rRNA ΔXᵢ comparison of two 30S replicates isolated from the cells expressing wild-type (WT) DbpA.** The ΔXᵢ is obtained from EpiNano as explained in Materials and Methods section of the manuscript. The nucleotides with no known modifications are shown in silver color, the nucleotides within five nucleotides of a known modification in gray, Ψ nucleotides are shown as red, m^7^G in magenta, m^2^G in dark purple, m^5^C in cyan, m^3^U in salmon pink, m^2^_6_A in gold, m^4^C_m_ in light green.

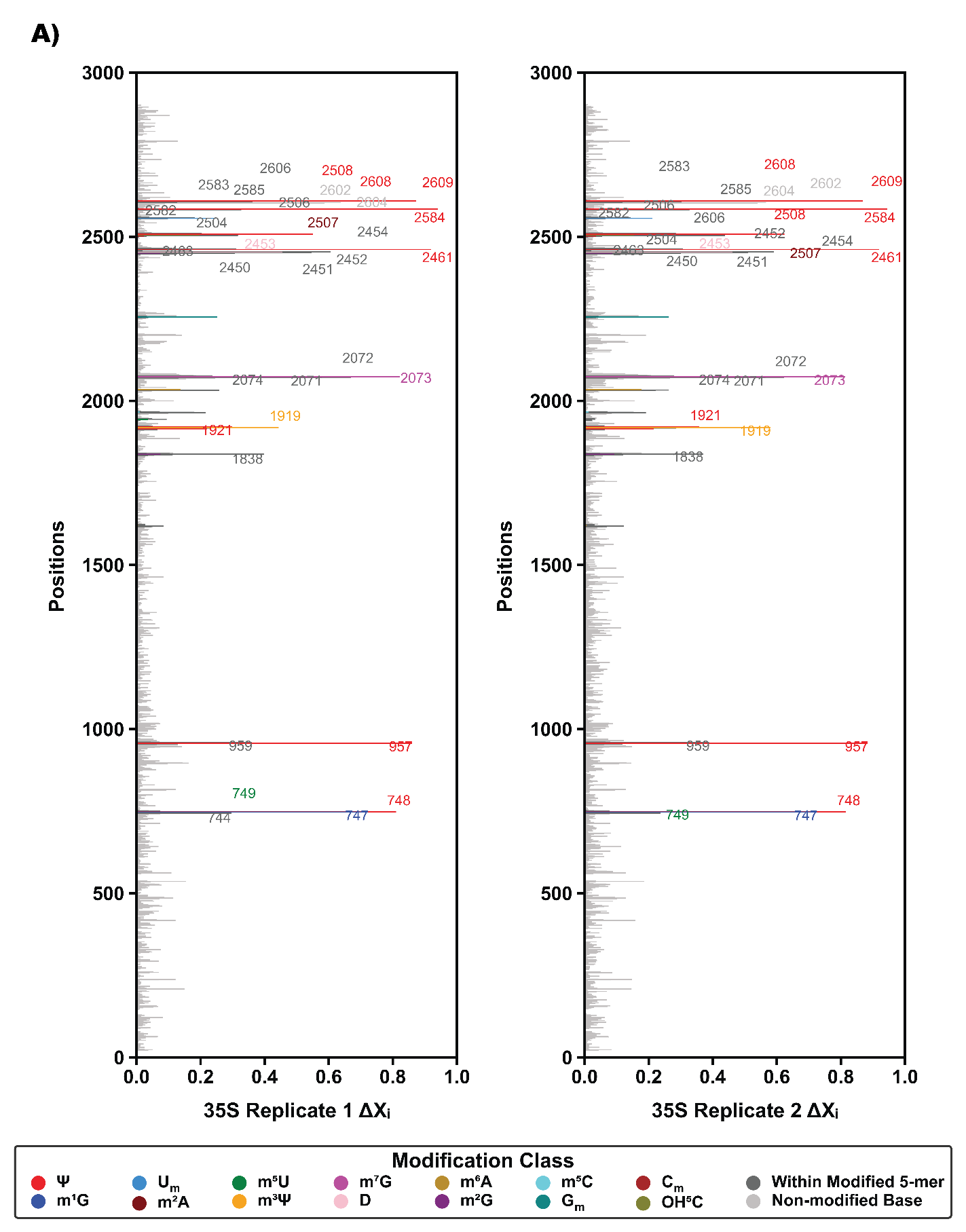

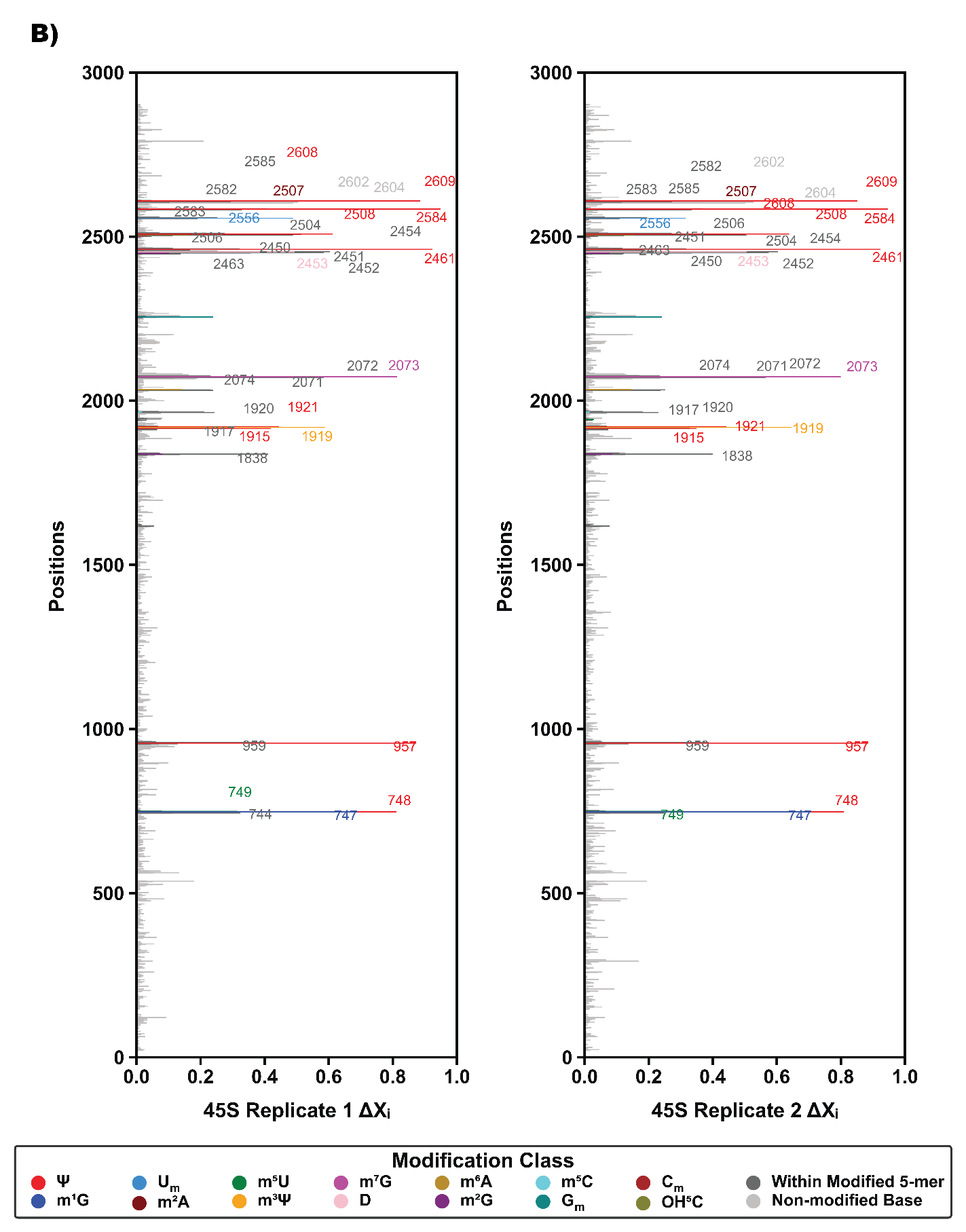

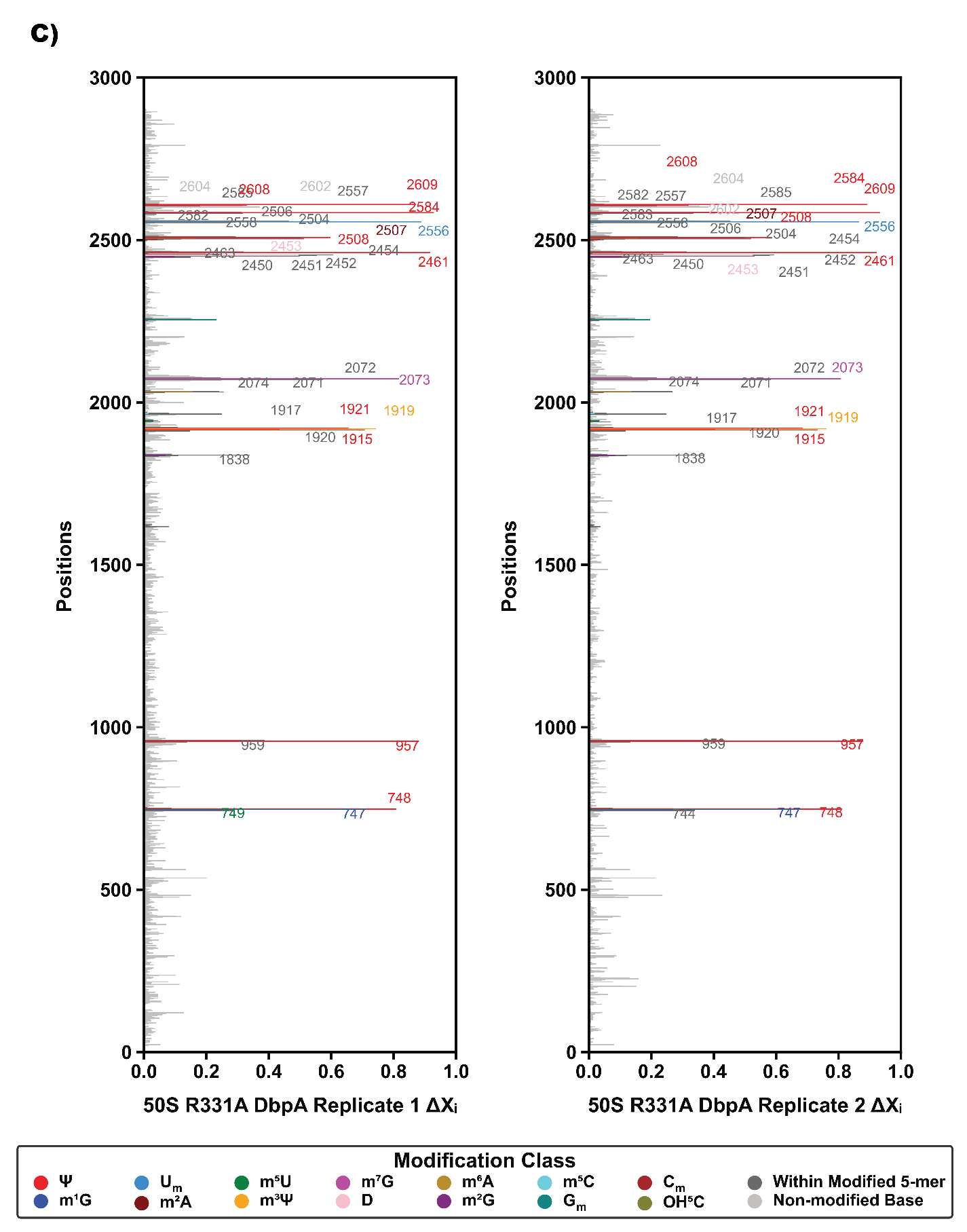

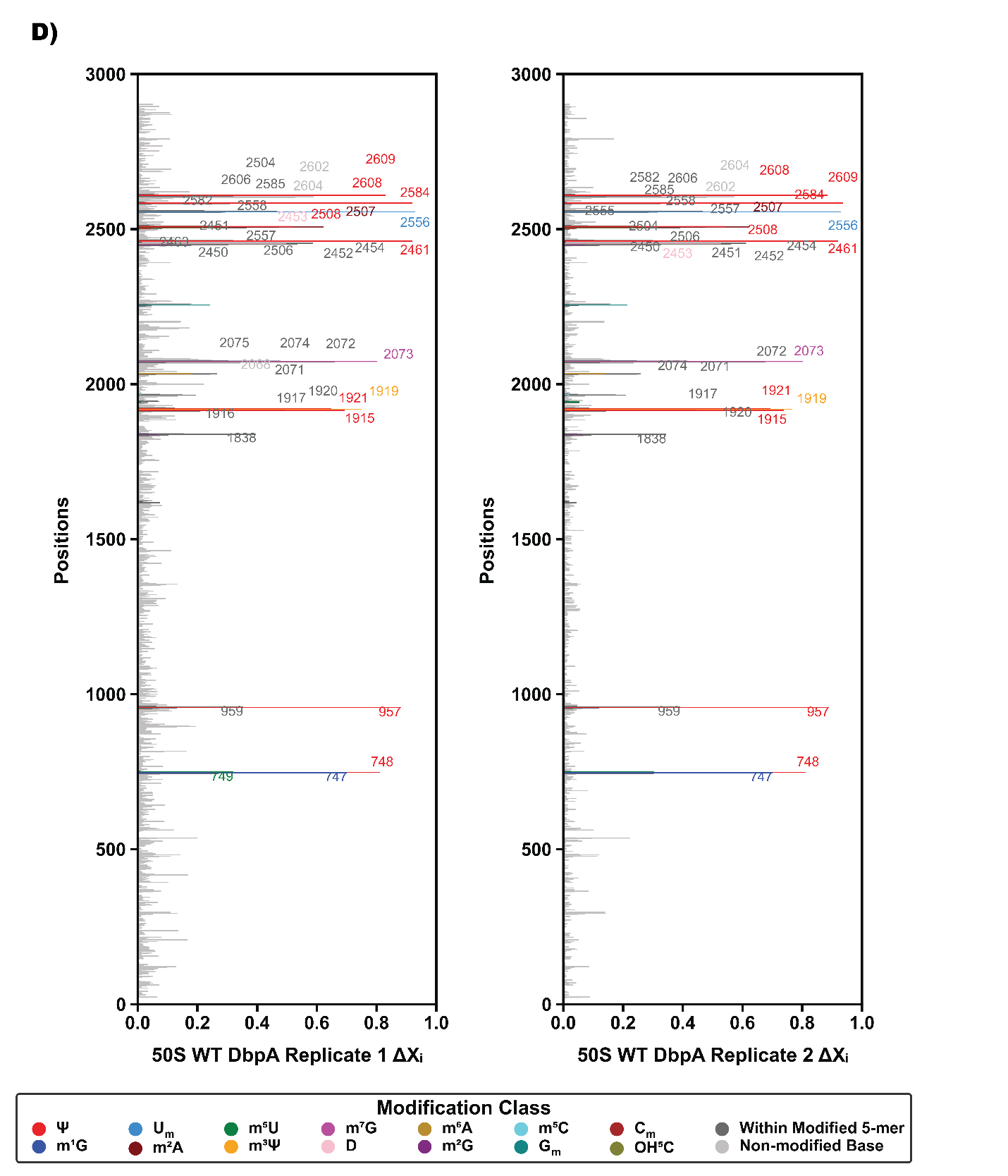

**Figure S6. ΔXᵢ of each nucleotide in both biological replicates of 35S (A), 45S (B), 50S accumulated in cells expressing R331A DbpA (C), 50S accumulated in cells expressing wild-type (WT) DbpA (D).** For all the panels in this figure, the Y axis is the 23S rRNA nucleotide position, the X axis is the ΔXᵢ, which was calculated from EpiNano as explained in the Materials and Methods section of the manuscript *^3^*. The nucleotides with no known modifications are shown in silver color lines, the nucleotides within five nucleotides of a known modification are shown as gray, the Ψ are shown as red, m^1^G as dark blue, U_m_ as sky blue, m^2^A as maroon, m^5^U as green, m^3^Ψ as orange, m^7^G as magenta, D as pink, m^6^A as golden, m^2^G as purple, m^5^C as cyan, G_m_ as teal, C_m_ as dark red, and OH^5^C as olive.

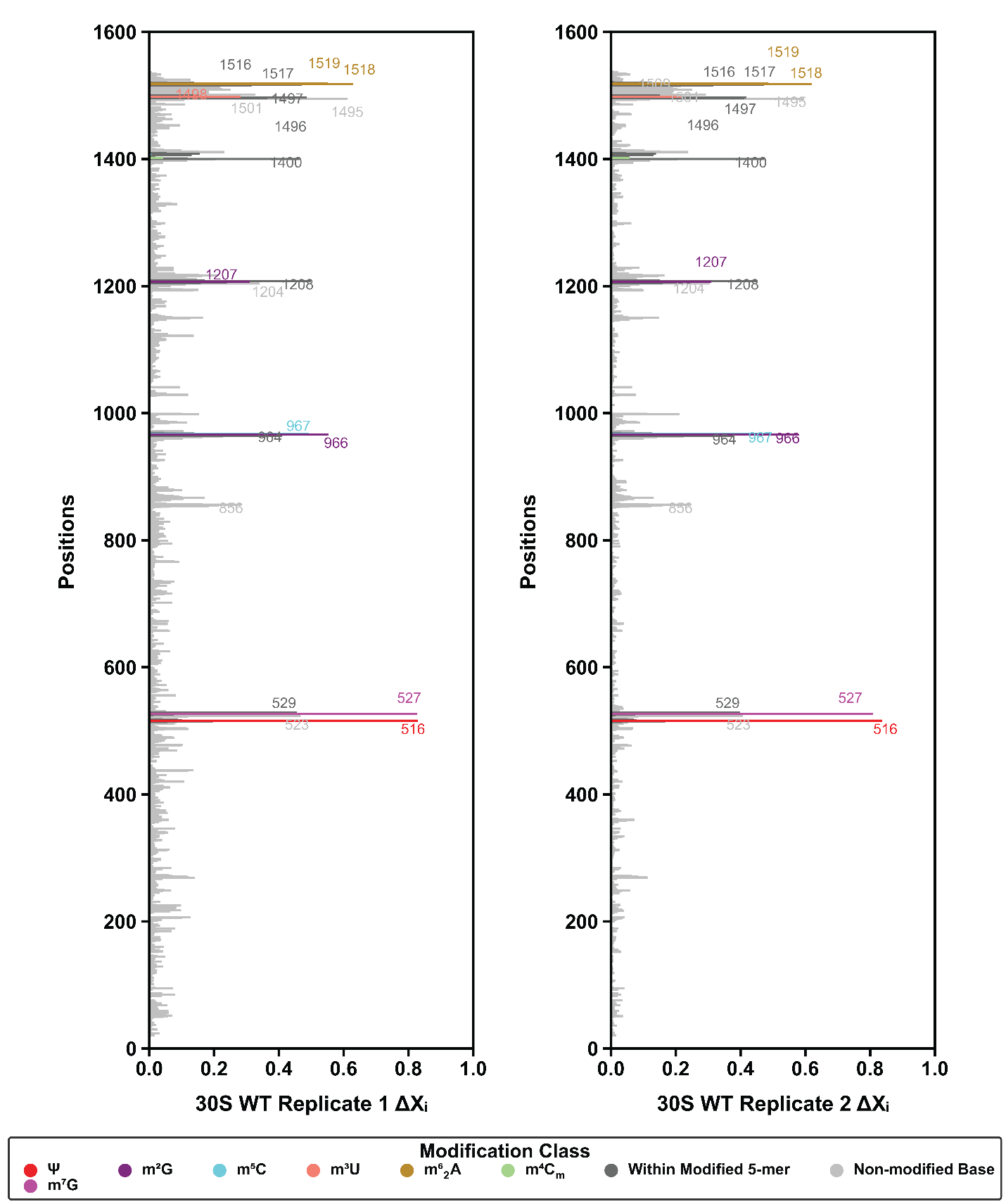

**Figure S7. ΔXᵢ of each nucleotide in both biological replicates of 30S small subunit from the cells expressing wild-type (WT) DbpA protein.** For all the panels in this figure the Y axis is the 16S rRNA nucleotide position, the X axis is the ΔXᵢ, which was calculated from EpiNano as explained in the Materials and Method section of the manuscript *^3^*. The nucleotides with no known modifications are shown in silver color lines, the nucleotides within a five nucleotide of a known modification are shown in gray, Ψ nucleotides are shown as red, m^7^G in magenta, m^2^G in dark purple, m^5^C in cyan, m^3^U in salmon pink, m^6^_2_A in gold, m^4^C_m_ in light green.

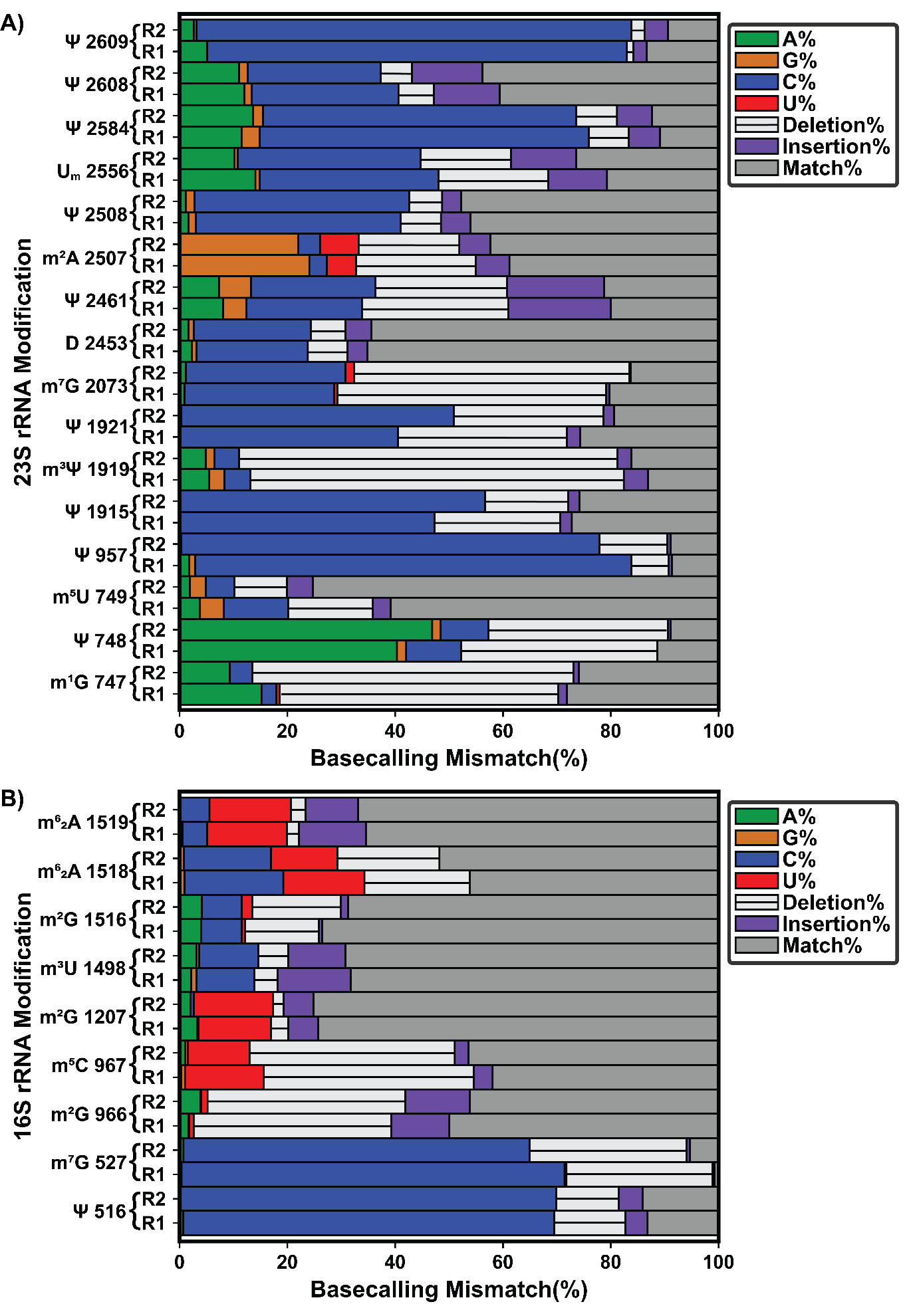

**Figure S8. Basecalling error profiles for RNA modifications in the 30S and 50S particles accumulated in cells expressing wild-type DbpA.** A) The error profiles of the modifications deemed detectable by our thresholds in the 23S rRNA from the 50S isolated from cells expressing wild-type DbpA. B) The error profiles of the modifications deemed detectable by our thresholds in the 16S rRNA from the 30S isolated from cells expressing wild-type DbpA. In both panels (A) and (B), fractions of mismatches, deletions, and insertions, for every base were calculated by dividing the number of times a mismatch, a deletion or insertion event occurred at a position by the position’s total read depth. A green, blue, orange, or red portion of a bar represents the fraction of the total read depth of a certain nucleotide that was miscalled as an A, C, G, or U, respectively.  A light gray bar with a black line in the middle of the bar represents the fraction of deletions at a specific position. A purple bar represents the percentage of insertions. In both panels (A) and (B), the error profiles of two biological replicates are shown. Biological replicate 1 is labeled as R1, while the biological replicate 2 is labeled as R2.

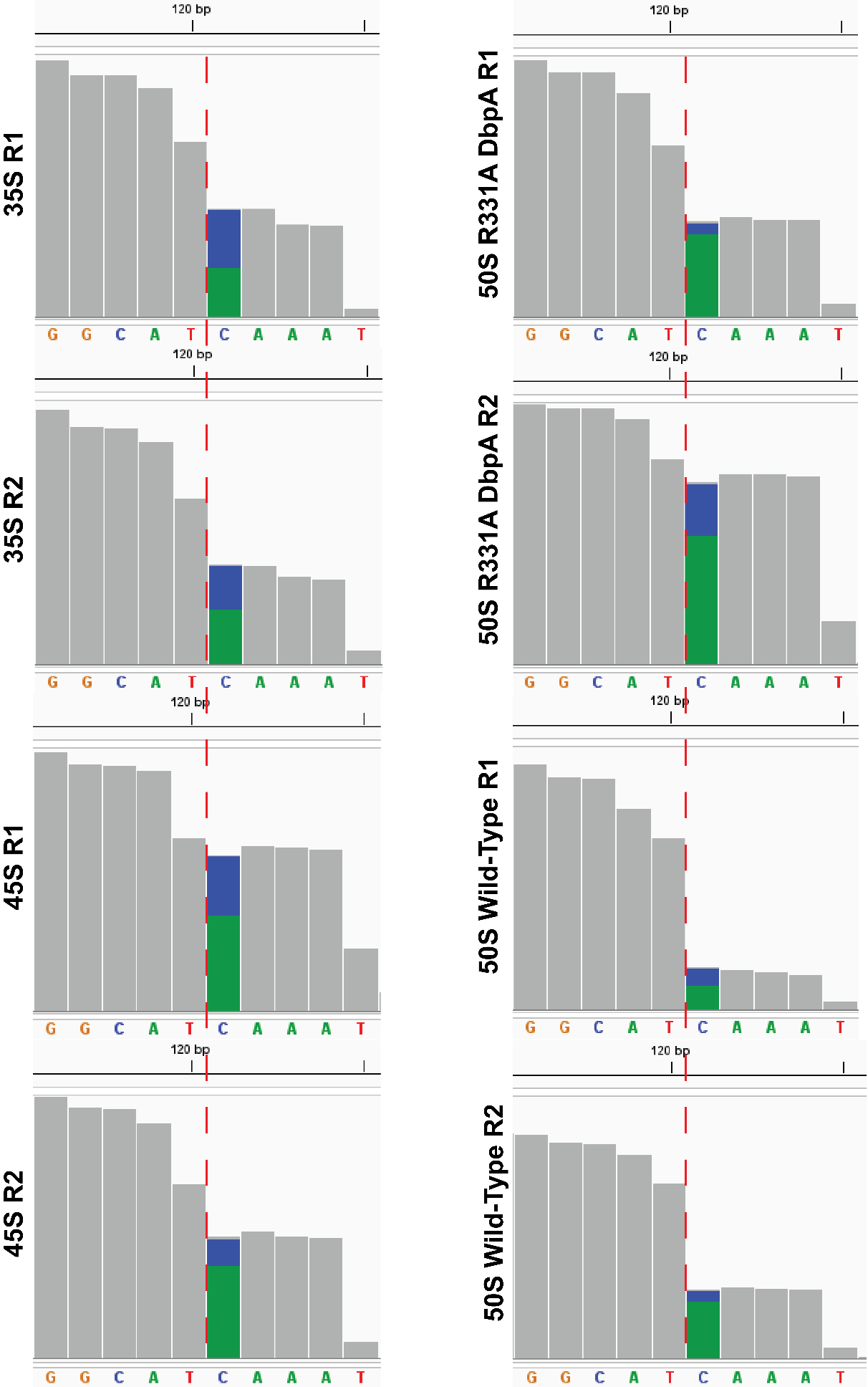

**Figure S9. 3’ end read depth of mature and unprocessed 5S rRNA in 35S, 45S, 50S particles from R331A DbpA, and 50S from wild-type DbpA cells.** The IGV snapshots of the sequence alignment to the *rrnB* gene of 5S rRNA from the 35S, 45S, 50S isolated from cells expressing R331A DbpA protein, and the 50S isolated from cells expressing wild-type DbpA. The figure shows the last 5 bases of mature 5S rRNA and the five extra nucleotides present in unprocessed 5S rRNA. The last position in the correctly matured 5S rRNA, residue 120, is marked with a red dotted line. R1 and R2 stand for biological replicates 1 and 2.

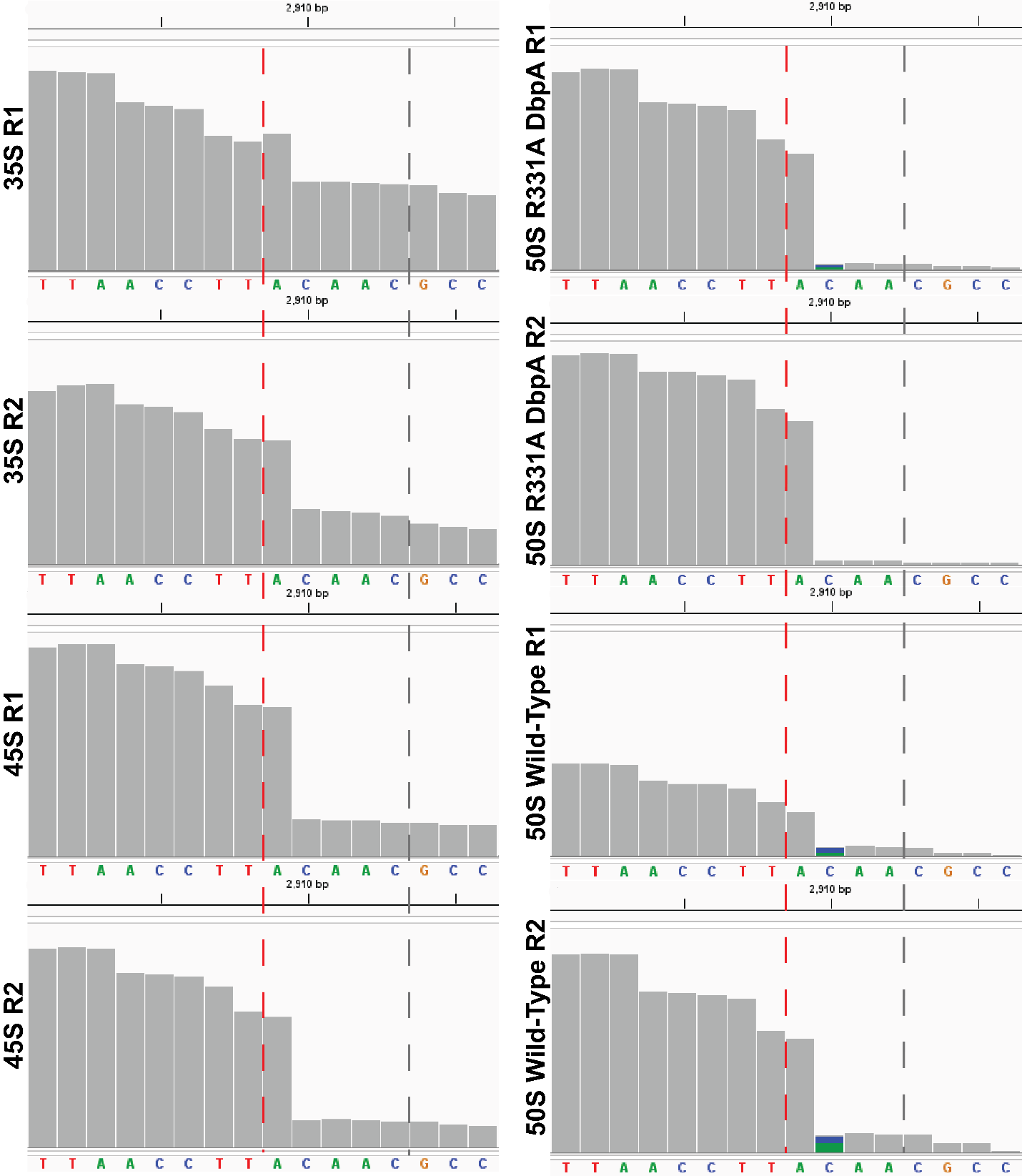

**Figure S10. 3’ end read depth of mature and unprocessed 23S rRNA in the 35S, 45S, 50S particles from R331A DbpA expressing cells, and the 50S from wild-type DbpA expressing cells.** The IGV snapshots from the sequence alignment to the *rrlB* gene of 23S rRNA from the 35S, 45S, 50S isolated from cells expressing R331A DbpA, and 50S isolated from cells expressing wild-type DbpA. Only the last 8 bases of mature 23S rRNA, and the 8 extra nucleotides present in unprocessed 23S rRNA are shown. The last position of properly matured 23S rRNA, residue 2908, is marked with a red dotted line, while the immature 23S rRNA containing five extra bases, is marked with a gray dotted line. R1 and R2 stand for biological replicates 1 and 2.

### **Table S1. List of modifications and the enzymes that incorporate the modifications in 23S rRNA and 16S rRNA of *E. coli*.**

| 23S rRNA Position^a^ | **Modification**^b^ | Enzyme^c^ |
| --- | --- | --- |
| 747 | **m^1^G** | RlmA*^4^* |
| 748 | **Ψ** | RluA*^4^* |
| 749 | **m^5^U** | RlmC*^4^* |
| 957 | **Ψ** | RluC*^4^* |
| 1620 | **m^6^A** | RlmF*^5^* |
| 1837 | **m^2^G** | RlmG*^6^* |
| 1915 | **Ψ** | RluD*^4^* |
| 1919 | **m³Ψ**^d^ | RluD, RlmH*^4, 7^* |
| 1921 | **Ψ** | RluD*^4^* |
| 1943 | **m⁵U** | RlmD*^4^* |
| 1966 | **m⁵C** | RlmI*^8^* |
| 2034 | **m⁶A** | RlmJ*^9^* |
| 2073 | **m⁷G** | RlmKL*^10^* |
| 2255 | **G_m_** | RlmB*^4^* |
| 2449 | **m²G** | RlmKL*^10^* |
| 2453 | **D^5S^_m_ ^e^** | RdsA, RlmX*^11, 12^* |
| 2461 | **Ψ** | RluE*^4^* |
| 2502 | **C_m_^5S^_m_^f^** | RlmM, RlmX*^12, 13^* |
| 2505 | **OH⁵C** | RlhA*^14^* |
| 2507 | **m²A** | RlmN*^15^* |
| 2508 | **Ψ** | RluC*^4^* |
| 2556 | **Um** | RlmE*^4^* |
| 2584 | **Ψ** | RluC*^4^* |
| 2608 | **Ψ** | RluF*^4^* |
| 2609 | **Ψ** | RluB*^4^* |
| 16S rRNA Position^a^ | **Modification** | Enzyme |
| 516 | **Ψ** | RsuA*^4^* |
| 527 | **m^7^G** | RsmG*^16^* |
| 966 | **m^2^G** | RsmD*^17^* |
| 967 | **m^5^C** | RsmB*^4^* |
| 1207 | **m^2^G** | RsmC*^4^* |
| 1402 | **m^4^C_m_**^g^ | RsmH, RsmI^e^*^18^* |
| 1407 | **m^5^C** | RsmF*^19^* |
| 1498 | **m^3^U** | RsmE*^20^* |
| 1516 | **m^2^G** | RsmJ*^21^* |
| 1518 | **m^6^_2_A** | RsmA*^4^* |
| 1519 | **m^6^_2_A** | RsmA*^4^* |

^a^ Positions of rRNA modifications are those corresponding to *rrlB* and *rrsB* genes that encode 23S rRNA and 16S rRNA, respectively.

^b^ Modifications found in the rRNA

^c^ The enzymes that incorporate the modification in the rRNA

^d^ RluD converts U 1919 of 23 rRNA to Ψ**,** subsequently RlmH methylates Ψ 1919 at position 3.*^4, 7^*

^e^ RdsA converts U 2453 of 23S rRNA to D, under anaerobic conditions, RlmX methylates 5’ carbon of D 2453.*^11, 12^*

^f^ RlmM methylates C 2502 of 23S rRNA at 2’-OH position, under anaerobic conditions, RlmX methylates 5’ carbon of C 2502. *^12, 13^*

^g^ RsmH and RsmL are responsible for the methylations at the N4 and 2’-OH positions of C1402 in the 16S rRNA, respectively.*^18^*

### **Table S2. Z-Score and SE fold change for large subunit particles isolated from cells expressing either the wild-type DbpA or R331A construct.**

| Position^a^ | Modification^b^ | Z score^c^ | SE Fold Change^c^ | Particle^d^ |
| --- | --- | --- | --- | --- |
| 747 | **m¹G** | 7.83 ± 0.02 | 4.07 ± 0.0 | 35S R331A |
| 748 | **Ψ** | 8.84 ± 0.01 | 2.99 ± 0.0 |  |
| 749 | **m⁵U** | 3.24 ± 0.04 | 3.4 ± 0.02 |  |
| 957 | **Ψ** | 9.51 ± 0.1 | 5.94 ± 0.02 |  |
| 1915 | **Ψ** | 2.09 ± 0.05 | 2.12 ± 0.01 |  |
| 1919 | **m³Ψ** | 5.42 ± 0.74 | 2.87 ± 0.17 |  |
| 1921 | **Ψ** | 3.34 ± 0.31 | 1.54 ± 0.09 |  |
| 2073 | **m⁷G** | 8.89 ± 0.09 | 5.75 ± 0.01 |  |
| 2453 | **D** | 4.82 ± 0.0 | 2.49 ± 0.01 |  |
| 2461 | **Ψ** | 10.05 ± 0.05 | 4.56 ± 0.0 |  |
| 2507 | **m²A** | 5.14 ± 0.05 | 4.36 ± 0.02 |  |
| 2508 | **Ψ** | 6.29 ± 0.37 | 3.05 ± 0.08 |  |
| 2556 | **U_m_** | 2.22 ± 0.23 | 2.13 ± 0.09 |  |
| 2584 | **Ψ** | 10.31 ± 0.03 | 4.48 ± 0.0 |  |
| 2608 | **Ψ** | 6.02 ± 0.38 | 2.81 ± 0.07 |  |
| 2609 | **Ψ** | 9.49 ± 0.07 | 4.07 ± 0.0 |  |
| 747 | **m¹G** | 7.47 ± 0.12 | 4.01 ± 0.02 | 45S R331A |
| 748 | **Ψ** | 8.73 ± 0.01 | 2.98 ± 0.0 |  |
| 749 | **m⁵U** | 3.16 ± 0.03 | 3.37 ± 0.01 |  |
| 957 | **Ψ** | 9.5 ± 0.09 | 5.95 ± 0.01 |  |
| 1915 | **Ψ** | 3.97 ± 0.38 | 2.79 ± 0.11 |  |
| 1919 | **m³Ψ** | 6.57 ± 0.34 | 3.1 ± 0.06 |  |
| 1921 | **Ψ** | 4.63 ± 0.01 | 1.84 ± 0.0 |  |
| 2073 | **m⁷G** | 8.68 ± 0.06 | 5.73 ± 0.01 |  |
| 2453 | **D** | 5.26 ± 0.08 | 2.6 ± 0.02 |  |
| 2461 | **Ψ** | 9.98 ± 0.02 | 4.56 ± 0.0 |  |
| 2507 | **m²A** | 5.36 ± 0.05 | 4.42 ± 0.02 |  |
| 2508 | **Ψ** | 6.66 ± 0.16 | 3.13 ± 0.03 |  |
| 2556 | **U_m_** | 4.17 ± 0.95 | 2.79 ± 0.27 |  |
| 2584 | **Ψ** | 10.26 ± 0.02 | 4.48 ± 0.0 |  |
| 2608 | **Ψ** | 5.01 ± 0.42 | 2.6 ± 0.09 |  |
| 2609 | **Ψ** | 9.38 ± 0.16 | 4.06 ± 0.03 |  |
| 747 | **m¹G** | 7.15 ± 0.21 | 4.01 ± 0.03 | 50S R331A |
| 748 | **Ψ** | 8.41 ± 0.02 | 2.98 ± 0.0 |  |
| 749 | **m⁵U** | 3.0 ± 0.15 | 3.35 ± 0.06 |  |
| 957 | **Ψ** | 9.17 ± 0.06 | 5.95 ± 0.0 | 50S R331A |
| 1915 | **Ψ** | 7.45 ± 0.11 | 3.6 ± 0.02 |  |
| 1919 | **m³Ψ** | 7.8 ± 0.06 | 3.36 ± 0.02 |  |
| 1921 | **Ψ** | 6.91 ± 0.13 | 2.29 ± 0.03 |  |
| 2073 | **m⁷G** | 8.45 ± 0.09 | 5.74 ± 0.01 |  |
| 2453 | **D** | 5.18 ± 0.15 | 2.63 ± 0.04 |  |
| 2461 | **Ψ** | 9.6 ± 0.02 | 4.56 ± 0.0 |  |
| 2507 | **m²A** | 5.06 ± 0.03 | 4.4 ± 0.01 |  |
| 2508 | **Ψ** | 6.06 ± 0.08 | 3.06 ± 0.01 |  |
| 2556 | **U_m_** | 9.15 ± 0.16 | 3.79 ± 0.02 |  |
| 2584 | **Ψ** | 9.72 ± 0.03 | 4.46 ± 0.0 |  |
| 2608 | **Ψ** | 3.17 ± 0.05 | 2.15 ± 0.02 |  |
| 2609 | **Ψ** | 9.19 ± 0.07 | 4.08 ± 0.02 |  |
| 747 | **m¹G** | 7.14 ± 0.08 | 4.01 ± 0.01 | 50S Wild-type |
| 748 | **Ψ** | 8.26 ± 0.04 | 2.97 ± 0.02 |  |
| 749 | **m⁵U** | 2.88 ± 0.12 | 3.33 ± 0.08 |  |
| 957 | **Ψ** | 9.16 ± 0.2 | 5.95 ± 0.0 |  |
| 1915 | **Ψ** | 7.24 ± 0.26 | 3.57 ± 0.02 |  |
| 1919 | **m³Ψ** | 7.76 ± 0.19 | 3.36 ± 0.0 |  |
| 1921 | **Ψ** | 6.73 ± 0.22 | 2.27 ± 0.01 |  |
| 2073 | **m⁷G** | 8.27 ± 0.16 | 5.71 ± 0.0 |  |
| 2453 | **D** | 4.48 ± 0.46 | 2.47 ± 0.14 |  |
| 2461 | **Ψ** | 9.05 ± 0.32 | 4.48 ± 0.07 |  |
| 2507 | **m²A** | 6.1 ± 0.1 | 4.65 ± 0.05 |  |
| 2508 | **Ψ** | 5.06 ± 0.08 | 2.85 ± 0.04 |  |
| 2556 | **U_m_** | 8.97 ± 0.49 | 3.78 ± 0.1 |  |
| 2584 | **Ψ** | 9.39 ± 0.0 | 4.42 ± 0.03 |  |
| 2608 | **Ψ** | 5.91 ± 0.41 | 2.84 ± 0.11 |  |
| 2609 | **Ψ** | 8.79 ± 0.35 | 4.03 ± 0.02 |  |

^a^ The position of the modifications in the 23S rRNA sequence.

^b^ The modifications shown are the ones deemed detectable using our thresholds (Materials and Methods)

^c^ The Z score and SE fold change were calculated as explained in the Materials and Methods section of the manuscript. The values shown for the Z-score and SE fold change are the average values obtained for two biological replicates, the errors are the standard deviations from the averages.

^d^ The 35S and 45S intermediates were isolated from R331A expressing cells (35S R331A, 45S R331A), the 50S particles were isolated either from wild-type DbpA expressing cells or R331A DbpA expressing cells (50S R331A)

### **Table S3. Z-Score and SE Fold Changes for the known modified positions present in the 16S rRNA of wild-type DbpA-expressing cells.**

| Position | Modification Class | Z-Score^b^ | SE Fold Change^b^ |
| --- | --- | --- | --- |
| **516** | **Ψ** | 10.85 ± 0.26 | 3.57 ± 0.01 |
| **527** | **m^7^G** | 10.65 ± 0.09 | 2.57 ± 0.01 |
| **966** | **m^2^G** | 7.24 ± 0.33 | 3.78 ± 0.03 |
| **967** | **m^5^C** | 6.23 ± 0.15 | 2.94 ± 0.0 |
| **1207** | **m^2^G** | 3.77 ± 0.12 | 3.93 ± 0.0 |
| **1402** | **m^4^C_m_** | 0.26 ± 0.17 | -0.47 ± 0.08 |
| **1407** | **m^5^C** | -0.38 ± 0.04 | 0.01 ± 0.01 |
| **1498** | **m^3^U** | 2.97 ± 0.31 | 1.21 ± 0.1 |
| **1516** | **m^2^G** | 3.85 ± 0.12 | 4.47 ± 0.0 |
| **1518** | **m^6^_2_A** | 8.03 ± 0.11 | 5.79 ± 0.01 |
| **1519** | **m^6^_2_A** | 6.59 ± 0.28 | 5.46 ± 0.09 |

^a^The position of the modifications in the 16S rRNA sequence. 16S rRNA was obtained from the 30S isolated from cells expressing the wild-type DbpA.

^b^ The Z score and SE Fold Change were calculated as explained in the Materials and Methods section of the manuscript. The values shown for the Z-score and SE fold change are the average values obtained for two biological replicates, the errors are the standard deviations from the averages.

### **Table S4. 95% confidence interval (CI) of Z-scores and Fold Change for all detected modifications in 23S rRNA and 16S rRNA.**

| **rRNA** | **rRNA Modification** | **95% CI** | | |
| --- | --- | --- | --- | --- |
|  |  | **Z-Score**^a^ | **SE Fold-Change**^a^ | **Particle**^b^ |
| 23S rRNA | **m¹G 747** | (7.800, 7.867) | (4.062, 4.069) | 35S R331A |
|  | **Ψ 748** | (8.818, 8.859) | (2.983, 2.993) |  |
|  | **m⁵U 749** | (3.185, 3.288) | (3.363, 3.431) |  |
|  | **Ψ 957** | (9.372, 9.642) | (5.909, 5.965) |  |
|  | **Ψ 1915** | (2.022, 2.159) | (2.109, 2.135) |  |
|  | **m³Ψ 1919** | (4.404, 6.445) | (2.628, 3.089) |  |
|  | **Ψ 1921** | (2.916, 3.766) | (1.416, 1.654) |  |
|  | **m⁷G 2073** | (8.761, 9.018) | (5.736, 5.758) |  |
|  | **D 2453** | (4.814, 4.817) | (2.484, 2.505) |  |
|  | **Ψ 2461** | (9.975, 10.12) | (4.554, 4.557) |  |
|  | **m²A 2507** | (5.075, 5.207) | (4.333, 4.389) |  |
|  | **Ψ 2508** | (5.770, 6.802) | (2.936, 3.152) |  |
|  | **U_m_ 2556** | (1.907, 2.539) | (2.005, 2.241) |  |
|  | **Ψ 2584** | (10.276, 10.347) | (4.472, 4.478) |  |
|  | **Ψ 2608** | (5.503, 6.544) | (2.714, 2.897) |  |
|  | **Ψ 2609** | (9.400, 9.586) | (4.061, 4.071) |  |
|  | **m¹G 747** | (7.297, 7.635) | (3.990, 4.040) | 45S R331A |
|  | **Ψ 748** | (8.711, 8.746) | (2.983, 2.987) |  |
|  | **m⁵U 749** | (3.118, 3.200) | (3.349, 3.388) |  |
|  | **Ψ 957** | (9.371, 9.628) | (5.937, 5.964) |  |
|  | **Ψ 1915** | (3.445, 4.504) | (2.633, 2.946) |  |
|  | **m³Ψ 1919** | (6.099, 7.034) | (3.020, 3.185) |  |
|  | **Ψ 1921** | (4.620, 4.641) | (1.832, 1.843) |  |
|  | **m⁷G 2073** | (8.603, 8.763) | (5.711, 5.745) |  |
|  | **D 2453** | (5.140, 5.372) | (2.577, 2.619) |  |
|  | **Ψ 2461** | (9.949, 10.014) | (4.561, 4.561) |  |
|  | **m²A 2507** | (5.284, 5.432) | (4.400, 4.445) |  |
|  | **Ψ 2508** | (6.442, 6.887) | (3.089, 3.162) |  |
|  | **U_m_ 2556** | (2.848, 5.492) | (2.392, 3.135) |  |
|  | **Ψ 2584** | (10.229, 10.291) | (4.482, 4.483) |  |
|  | **Ψ 2608** | (4.426, 5.588) | (2.471, 2.727) |  |
|  | **Ψ 2609** | (9.149, 9.606) | (4.027, 4.100) |  |
|  | **m¹G 747** | (6.862, 7.445) | (3.960, 4.051) | 50S R331A |
|  | **Ψ 748** | (8.378, 8.441) | (2.979, 2.986) |  |
|  | **m⁵U 749** | (2.793, 3.199) | (3.272, 3.435) |  |
|  | **Ψ 957** | (9.091, 9.252) | (5.948, 5.953) |  |
|  | **Ψ 1915** | (7.295, 7.605) | (3.565, 3.632) |  |
| 23S rRNA | **m³Ψ 1919** | (7.712, 7.882) | (3.339, 3.381) | 50S R331A |
|  | **Ψ 1921** | (6.728, 7.101) | (2.256, 2.326) |  |
|  | **m⁷G 2073** | (8.324, 8.568) | (5.727, 5.750) |  |
|  | **D 2453** | (4.974, 5.389) | (2.575, 2.674) |  |
|  | **Ψ 2461** | (9.572, 9.634) | (4.551, 4.561) |  |
|  | **m²A 2507** | (5.017, 5.107) | (4.390, 4.404) |  |
|  | **Ψ 2508** | (5.952, 6.175) | (3.039, 3.073) |  |
|  | **U_m_ 2556** | (8.922, 9.374) | (3.771, 3.818) |  |
|  | **Ψ 2584** | (9.685, 9.760) | (4.453, 4.462) |  |
|  | **Ψ 2608** | (3.098, 3.238) | (2.129, 2.176) |  |
|  | **Ψ 2609** | (9.086, 9.292) | (4.059, 4.106) |  |
|  | **m¹G 747** | (7.028, 7.260) | (3.991, 4.028) | 50S Wild-type |
|  | **Ψ 748** | (8.201, 8.319) | (2.938, 2.992) |  |
|  | **m⁵U 749** | (2.719, 3.048) | (3.216, 3.432) |  |
|  | **Ψ 957** | (8.888, 9.441) | (5.952, 5.954) |  |
|  | **Ψ 1915** | (6.872, 7.599) | (3.541, 3.590) |  |
|  | **m³Ψ 1919** | (7.493, 8.021) | (3.353, 3.363) |  |
|  | **Ψ 1921** | (6.420, 7.043) | (2.249, 2.283) |  |
|  | **m⁷G 2073** | (8.041, 8.494) | (5.711, 5.717) |  |
|  | **D 2453** | (3.848, 5.120) | (2.276, 2.658) |  |
|  | **Ψ 2461** | (8.612, 9.493) | (4.377, 4.584) |  |
|  | **m²A 2507** | (5.970, 6.235) | (4.578, 4.716) |  |
|  | **Ψ 2508** | (4.956, 5.168) | (2.788, 2.911) |  |
|  | **U_m_ 2556** | (8.297, 9.645) | (3.638, 3.906) |  |
|  | **Ψ 2584** | (9.385, 9.396) | (4.375, 4.455) |  |
|  | **Ψ 2608** | (5.336, 6.483) | (2.690, 2.990) |  |
|  | **Ψ 2609** | (8.303, 9.274) | (3.994, 4.061) |  |
| 16S rRNA | **Ψ 516** | (10.482, 11.216) | (2.553, 2.589) | 30S Wild-type |
|  | **m⁷G 527** | (10.529, 10.765) | (3.736, 3.819) |  |
|  | **m²G 966** | (6.780, 7.708) | (2.942, 2.944) |  |
|  | **m⁵C 967** | (6.021, 6.447) | (3.930, 3.934) |  |
|  | **m²G 1207** | (3.601, 3.941) | (1.066, 1.349) |  |
|  | **m³U 1498** | (2.532, 3.404) | (4.466, 4.466) |  |
|  | **m²G 1516** | (3.683, 4.013) | (5.775, 5.805) |  |
|  | **m⁶₂A 1518** | (7.881, 8.181) | (5.339, 5.581) |  |
|  | **m⁶₂A 1519** | (6.199, 6.973) | (2.553, 2.589) |  |

^a^ The Z-score and SE Fold Change values were calculated as explained in the Materials and Methods section of the manuscript. The 95% CI was calculated using the equation below.

$CI=\bar{x} \pm1.96 \frac{std}{\sqrt{2}}$	Equation 1

In Equation 1, $\bar{x}$ is the mean of Z-score or Fold Change between the biological replicates; std refers to the standard deviation between the biological replicates, and n is the sample size, which is 2.

^b^ Particles were isolated from cells expressing either the R331A or wild-type DbpA protein as explained in the Materials and Methods section of the manuscript.

### **REFERENCES**

[1] Robinson, J. T., Thorvaldsdóttir, H., Winckler, W., Guttman, M., Lander, E. S., Getz, G., and Mesirov, J. P. (2011) Integrative genomics viewer, *Nature Biotechnology* *29*, 24-26.

[2] Li, H. (2018) Minimap2: pairwise alignment for nucleotide sequences, *Bioinformatics* *34*, 3094-3100.

[3] Liu, H., Begik, O., Lucas, M. C., Ramirez, J. M., Mason, C. E., Wiener, D., Schwartz, S., Mattick, J. S., Smith, M. A., and Novoa, E. M. (2019) Accurate detection of m(6)A RNA modifications in native RNA sequences, *Nat Commun* *10*, 4079.

[4] Ofengand, J., and Campo, M. D. (2004-12-29) Modified Nucleosides of Escherichia coli Ribosomal RNA, *EcoSal Plus* *1*.

[5] Sergiev, P. V., Serebryakova, M. V., Bogdanov, A. A., and Dontsova, O. A. (2008) The ybiN Gene of Escherichia coli Encodes Adenine-N6 Methyltransferase Specific for Modification of A1618 of 23 S Ribosomal RNA, a Methylated Residue Located Close to the Ribosomal Exit Tunnel, *Journal of Molecular Biology* *375*, 291-300.

[6] Sergiev, P. V., Lesnyak, D. V., Bogdanov, A. A., and Dontsova, O. A. (2006) Identification of Escherichia coli m2G methyltransferases: II. The ygjO Gene Encodes a Methyltransferase Specific for G1835 of the 23 S rRNA, *Journal of Molecular Biology* *364*, 26-31.

[7] Ero, R., Peil, L., Liiv, A., and Remme, J. (2008) Identification of pseudouridine methyltransferase in <i>Escherichia coli</i>, *RNA* *14*, 2223-2233.

[8] Purta, E., O’Connor, M., Bujnicki, J. M., and Douthwaite, S. (2008) YccW is the m5C Methyltransferase Specific for 23S rRNA Nucleotide 1962, *Journal of Molecular Biology* *383*, 641-651.

[9] Golovina, A. Y., Dzama, M. M., Osterman, I. A., Sergiev, P. V., Serebryakova, M. V., Bogdanov, A. A., and Dontsova, O. A. (2012) The last rRNA methyltransferase of <i>E. coli</i> revealed: The <i>yhiR</i> gene encodes adenine-N6 methyltransferase specific for modification of A2030 of 23S ribosomal RNA, *RNA* *18*, 1725-1734.

[10] Wang, K.-T., Desmolaize, B., Nan, J., Zhang, X.-W., Li, L.-F., Douthwaite, S., and Su, X.-D. (2012) Structure of the bifunctional methyltransferase YcbY (RlmKL) that adds the m 7 G2069 and m 2 G2445 modifications in Escherichia coli 23S rRNA, *Nucleic Acids Research* *40*, 5138-5148.

[11] Toubdji, S., Thullier, Q., Kilz, L.-M., Marchand, V., Yuan, Y., Sudol, C., Goyenvalle, C., Jean-Jean, O., Rose, S., Douthwaite, S., Hardy, L., Baharoglu, Z., De Crécy-Lagard, V., Helm, M., Motorin, Y., Hamdane, D., and Brégeon, D. (2024) Exploring a unique class of flavoenzymes: Identification and biochemical characterization of ribosomal RNA dihydrouridine synthase, *Proceedings of the National Academy of Sciences* *121*.

[12] Ishiguro, K., Midorikawa, K., Shigi, N., Kimura, S., Liiv, A., Yokoyama, T., Ito, T., Shirouzu, M., Remme, J., Miyauchi, K., and Suzuki, T. (2025) Hypoxia-induced ribosomal RNA modifications in the peptidyl-transferase center contribute to anaerobic growth of bacteria, *Molecular Cell*.

[13] Purta, E., O'Connor, M., Bujnicki, J. M., and Douthwaite, S. (2009) YgdE is the 2′‐<i>O‐</i>ribose methyltransferase RlmM specific for nucleotide C2498 in bacterial 23S rRNA, *Molecular Microbiology* *72*, 1147-1158.

[14] Kimura, S., Sakai, Y., Ishiguro, K., and Suzuki, T. (2017) Biogenesis and iron-dependency of ribosomal RNA hydroxylation, *Nucleic Acids Research* *45*, 12974-12986.

[15] Toh, S.-M., Xiong, L., Bae, T., and Mankin, A. S. (2008) The methyltransferase YfgB/RlmN is responsible for modification of adenosine 2503 in 23S rRNA, *RNA* *14*, 98-106.

[16] Okamoto, S., Tamaru, A., Nakajima, C., Nishimura, K., Tanaka, Y., Tokuyama, S., Suzuki, Y., and Ochi, K. (2007) Loss of a conserved 7‐methylguanosine modification in 16S rRNA confers low‐level streptomycin resistance in bacteria, *Molecular Microbiology* *63*, 1096-1106.

[17] Lesnyak, D. V., Osipiuk, J., Skarina, T., Sergiev, P. V., Bogdanov, A. A., Edwards, A., Savchenko, A., Joachimiak, A., and Dontsova, O. A. (2007) Methyltransferase That Modifies Guanine 966 of the 16 S rRNA, *Journal of Biological Chemistry* *282*, 5880-5887.

[18] Kimura, S., and Suzuki, T. (2010) Fine-tuning of the ribosomal decoding center by conserved methyl-modifications in the Escherichia coli 16S rRNA, *Nucleic Acids Research* *38*, 1341-1352.

[19] Andersen, N. M., and Douthwaite, S. (2006) YebU is a m5C Methyltransferase Specific for 16 S rRNA Nucleotide 1407, *Journal of Molecular Biology* *359*, 777-786.

[20] Basturea, G. N., Rudd, K. E., and Deutscher, M. P. (2006) Identification and characterization of RsmE, the founding member of a new RNA base methyltransferase family, *RNA* *12*, 426-434.

[21] Basturea, G. N., Dague, D. R., Deutscher, M. P., and Rudd, K. E. (2012) YhiQ Is RsmJ, the Methyltransferase Responsible for Methylation of G1516 in 16S rRNA of E. coli, *Journal of Molecular Biology* *415*, 16-21.
